# 3D Focused Ion Beam Microscopy of Fossilized *Albertosaurus sarcophagus* reveals Nano to Microscale Structures

**DOI:** 10.1101/2025.04.13.648590

**Authors:** Alyssa Williams, Dirk Schumann, Jordan C. Mallon, Michael W. Phaneuf, Nabil Bassim, Kathryn Grandfield

**Author notes:** Corresponding author: Prof. Kathryn Grandfield McMaster University 1280 Main Street West Hamilton, L8S 4L7 Ontario, Canada.

## Abstract

Osteohistological investigations of fossilized bone can reveal details about the specimen’s biological, geological and environmental conditions. Micro-to-nanoscale imaging provides insight into the structural organization of bone and can also reveal indicators of the fossilization process. We examined a petrographic thin section of the left fibula of a ∼71.5 million-year-old *Albertosaurus sarcophagus* (Canadian Museum of Nature [CMN] catalogue number FV 11315) using nanoscale scanning electron microscopy (SEM) and focused ion beam (FIB)-SEM tomographic imaging to study the arrangement of mineral and organic components of fossil bone in multidimensions. Here, we present evidence of permineralization in Haversian canals by energy dispersive X-ray spectroscopy. Nanoscale 3D FIB-SEM imaging revealed that the characteristic 67 nm banding periodicity of collagen fibrils was remarkably well preserved over 70M years, and 3D imaging allowed for the detection of collagen fibril bundles in parallel fibered and lamellar bone arrangements. A newly discovered structure in modern bone, the ellipsoidal mineral cluster, was tiled throughout the 3D space of fibrolamellar fossil bone. These observations, afforded by the high-resolution and site-specific nature of FIB-SEM, link key fossilized features with the micro-nanoscale structure of modern-day bone. This investigation highlights the persistence of bone formation and organization persisting for over millions of years.

## INTRODUCTION

Fossil osteohistological investigations detail the microstructural level of bone preservation and provide insight into the specimens’ greater environment and biology. The organization of the collagen and mineral system is key to the development of normal, unaltered bones in living animals. However, understanding this association in extinct, dead animals is challenging due to the fossilization process. Over time, fossilization and diagenetic processes change the structure and properties of bone^1–4^. Changes in fossil bone mineral content are key markers of diagenesis^1–3,5^. Carbonated hydroxyapatite, the inorganic phase original to bone, can undergo dissolution and recrystallization, resulting in changes to crystal size and lattice arrangement, as well as potential incorporation of trace minerals introduced from the environment^1–3,5^. Fossil bone is also subject to permineralization, where minerals dissolved in groundwater can infiltrate the bone and deposit in its pores and cavities^1,3,4^. Three processes of permineralization include pyritization, carbonate mineralization and silicification, which respectively lead to the deposition of pyrite, calcium carbonate and silica-based composites within bone^6–8^. After specimen death, soft tissue is subject to decay or preservation depending on the environment^4,9–11^. For example, rapid burial and anoxic environments can promote soft tissue preservation^4,12^. Soft tissue, such as skin^13^, muscle^14^, fibrous matrices^9,11,15–17^, and cells^9,11,17–20^ have been visualized in fossil bones with multiple imaging techniques. However, preserved soft tissue may undergo alterations in the structure of organic matter^21^. These diagenetic and fossilization processes lead to changes within the fossil bone, creating a denser and mineralogically complex material. Investigating the arrangement of collagen and mineral is key to understanding bone function, development processes and the preservation of the collagen fibril and mineral system in fossil bone.

An emergent nanoscale 3D imaging technique utilized to analyze bone in recent years is focused ion beam scanning electron microscopy (FIB-SEM), also referred to as FIB-SEM nanotomography. During FIB-SEM nanotomography, serial sectioning with an ion beam and imaging with an electron beam produces a 2D image stack of the probed volume that can be reconstructed and rendered in 3D to elucidates the nanoscale structural arrangement, or ultrastructure, of minerals and collagen in bone tissue. ^22–24^. Bone tissue has a hierarchical system from the skeletal to the molecular level contributing to the bone’s structure and function^25–27^. Bone can be described based on the arrangement and composition of its mineral and organic components^25,26,28,29^. The collagen fibril network, hierarchically expanding from molecule to fibrils with banding periodicity to fibre bundles, has been well-documented in multiple modern species^25–27,29–35^. Mineralization of the collagen fibril network in lamellar bone occurs from mineral foci in the collagenous matrix that grow into ellipsoidal mineral clusters, eventually leading to a tessellated mineralization pattern ^36–38^. Bone cells, including osteoblasts, osteoclasts and osteocytes, play a key role in regulating bone formation, remodelling and mechanosensing, respectively^39–47^. Osteocytes reside in the lacunocanalicular network (LCN) in bone, where the cell body sits in lacunae with its processes extending through small channels, canaliculi, terminating at other cells or Haversian canals, the central supply of oxygen and nutrients to bone^42,44–46^. The structural organization of bone and its components has been extensively studied using 2D and 3D characterizations with light, x-ray and electron microscopy, for example ^28,37,38,48–52^.

In rapidly growing animals, including mammals and juvenile dinosaurs, fibrolamellar bone which is a transient, composite bone tissue is characteristic^7,53–63^. Elements of woven, parallel-fibered and lamellar bone fibril arrangements can be found in the microstructure of fibrolamellar bone tissue^7,53,55,56,64,65^. The woven arrangement is characterized by randomly arranged collagen fibrils with low mechanical advantages and is associated with fast growth^56,64–67^. The parallel fibered arrangement is characterized by the parallel orientation of collagen fibrils and is referred to as an intermediate between lamellar bone and woven bone^27,56,62,64^. The lamellar arrangement is characterized by fibrils arranged in concentric lamellae around blood vessels ^64,68^. Fibrolamellar bone formation begins with a blood vessel network that guides the formation of new bone^64,69^. Woven bone is deposited around blood vessels, forming a framework for further new bone formation^56,64,65^. Parallel fibered bone can be deposited onto this scaffold or without the scaffold to develop the bone further^7,56,64^. Lamellar bone is then formed, filling in spaces from the woven and parallel-fibered bone to the blood vessels within the bone^56,62,64,68^. Primary osteons are a key characteristic of the fibrolamellar bone surrounding blood vessels within the bone that also join with neighbouring bone^64^. The composition of fibrolamellar bone tissue can evolve during the development of the specimen as mechanical and physiological requirements change during the lifespan^7,53,58,64^.

In this study, 2D and 3D electron microscopy imaging and spectroscopy tools are employed to investigate the micro and nanoscale features of *Albertosaurus sarcophagus (A. sarcophagus*) bone from the Horseshoe Canyon Formation to understand how bone developed and was conserved over time. *A. Sarcophagus* is a theropod dinosaur in the family Tyrannosauridae whose fossils definitively occur in the lower two-thirds of the Horseshoe Canyon Formation in Alberta, Canada^61,70–74^. Recent isotopic dating yields an age range of 73.1–68.0 Ma for these strata in the type area^75^. *A. Sarcophagus* is among the earliest- and best-known tyrannosaurids, represented by several complete skeletons and bonebed material^73^. Consequently, it has been the subject of ongoing investigation, particularly as concerns the behaviour^76^, biomechanics^77^, and growth^72,77–80^ of the species. By employing nanoscale imaging techniques, we can analyze the conserved and unique features of *A.sarcophagus* bone, providing greater insight into bone development and fossilization in the Horseshoe Canyon Formation.

## RESULTS AND DISCUSSION

### 1.1 Imaging Overview of the Albertosaurus sarcophagus (CMNFV 11315)

The late juvenile–early subadult tyrannosaurid, CMNFV 11315, was previously identified as *Albertosaurus sarcophagus* based on morphometric and cladistic analyses, and gross osteohistological analysis of the fibula has established a minimum age at the time of death of approximately 2 years^61^. However, in the present study, the micron and nanoscale-level osteohistological features of this specimen are analyzed further to understand the growth and mineralization processes that occurred pre- and post-mortem. Features of interest for chemical analysis and 3D nanotomography were identified from the mosaic imaging found in the Browser-Based viewer (BBV) data, where the EDS maps and movie clips of the 3D nanotomography have also been linked. Access data here: https://www.petapixelproject.com/mosaics/museumofnature/CMNFV-11315/index.html

Fibrolamellar bone is the primary tissue in this fibula-thin section, with approximately two annuli visible in the lateral periosteal region of the bone^61^. The annuli are not directly continuous on the medial side due to bone remodelling^61^. The cortex region displays a highly mineralized matrix with no medullary cavity present (Fig. 1A & C). The lateral region, in comparison, is less mineralized, and several primary osteons and the Volkmann’s canals can be visualized. The medial side displays more remodelling in contrast to the lateral side, with overlapping secondary osteons present (Fig. 1B, D, E, G). 2D and 3D analysis was primarily performed on the lateral side of the fibula thin section.

**Figure 1.**
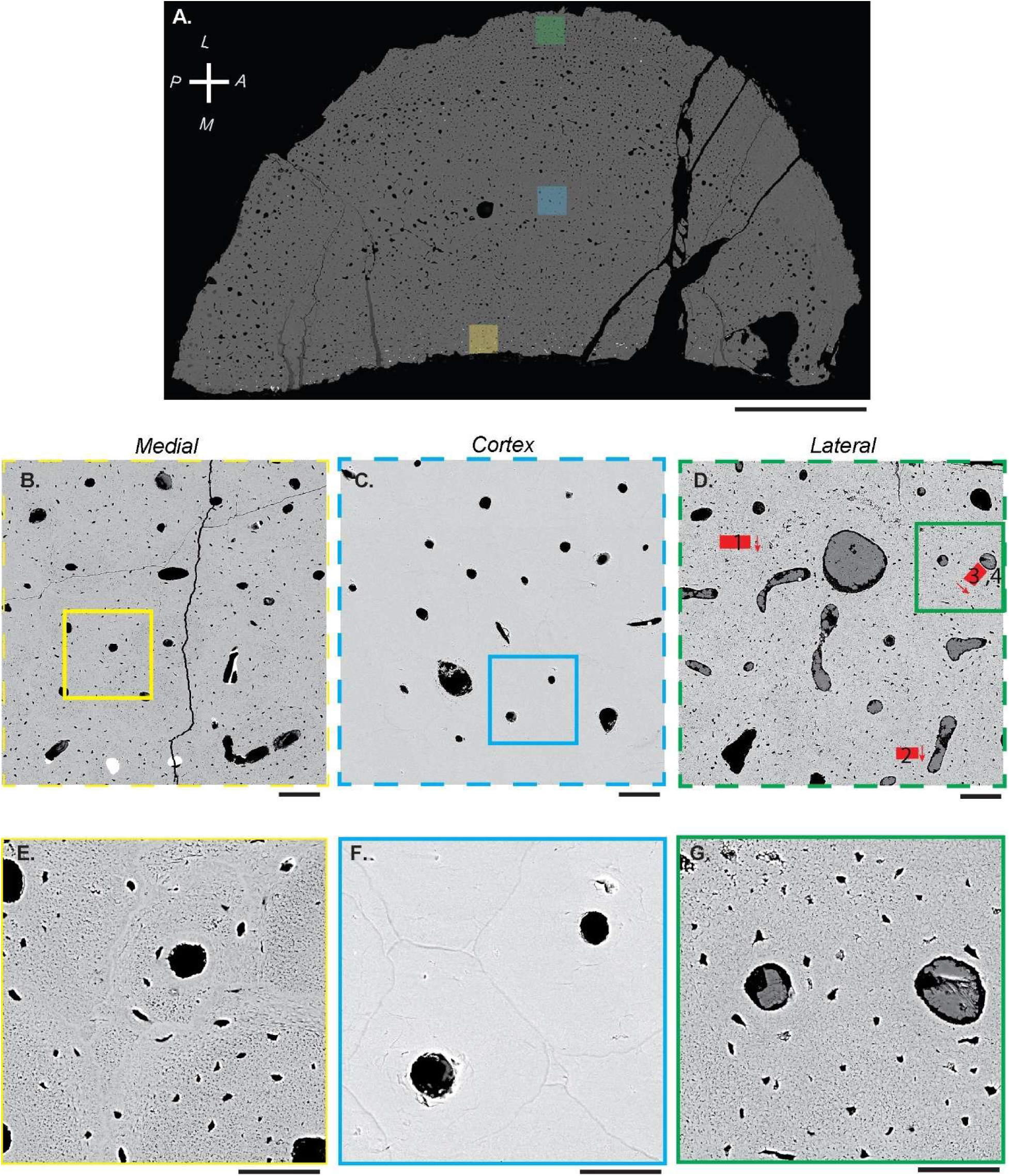
Backscattered electron images show an overview of the medial, cortex and lateral regions of the CMNFV 11315 D-shaped petrographic fibula thin section (Reproduced with permissions^61^). A) large-area mosaic image of the thin section with anatomical planes indicated. B) Medial region (yellow) of the fibula with overlapping secondary osteons. C) Cortex region (blue) of the fibula with highly mineralized bone matrix surrounding canals. D) Lateral fibula region (green) has several primary osteons and Volkmann’s canals present. E, F,G) Insets indicated in B, C, D, respectively, where E) overlapping secondary osteons with truncated cement lines are visible, F) highly mineralized bone matrix with mineralized lacunae surrounding the canal, and G) displays primary osteons without visible cement lines and dark lacunae. Scale bars A) 5 mm, B-D) 100 µm, F-G) 50 µm.

### 1.2 Elemental Analysis of Permineralization

Evidence of permineralization and other diagenetic processes was seen in the both the canals (Haversian and Volkmann’s) and osteocyte lacunocanalicular network (LCN) of the fossil after investigation with EDS (Figs. 2-4). Diagenetic processes, including cell death and fluid flow through the sediment formation, are facilitated by mineral dissolution and precipitation during fossilization. These diagenetic fluids can take advantage of the porosity connecting the vascular systems (Haversian and Volkmann’s) to the lacunocanalicular network (LCN), leading to secondary mineralization as was noted in lacunae (Fig. 2, S2), and in the Haversian canals in the lateral and medial regions of the fibula, which were determined to consist of calcite and baryte (Figs. 2A, B, F, H, S2, BBV dataset). Some canals were filled with kaolinite (Fig. 2C). Smectite-group minerals often lined the inner wall of the Haversian canals and filled interstices between other minerals (Fig. 2A-D,G, 3A, S2; also see BBV dataset). Pyrite occurs as beautifully formed framboidal pyrite, as sunflower framboidal pyrite grains (Fig 3, S1A-B) and as blocky pyrite crystals, which developed from precursor framboids (Figs. 2E, S1C-D and S2). Fluid flow during diageneses and burial also caused the dissolution of previously precipitated calcite (Fig. 2). The Haversian canals in the cortex region of the fibula showed fewer mineral fillings. Cracks in the fibula that likely were caused during burial and diagenesis filled with calcite (Fig. 2H). Similar stages of permineralization have been shown in *Metoposaurus* fossils occurring within a bonebed hosted by a mudstone layer^81^. In these *Metoposaurus* fossils, bacterial activity and clay mineral uptake occur initially, followed by pyrite and granular sparite, and then finally, barite and blocky sparite^81^.

**Fig. 2.**
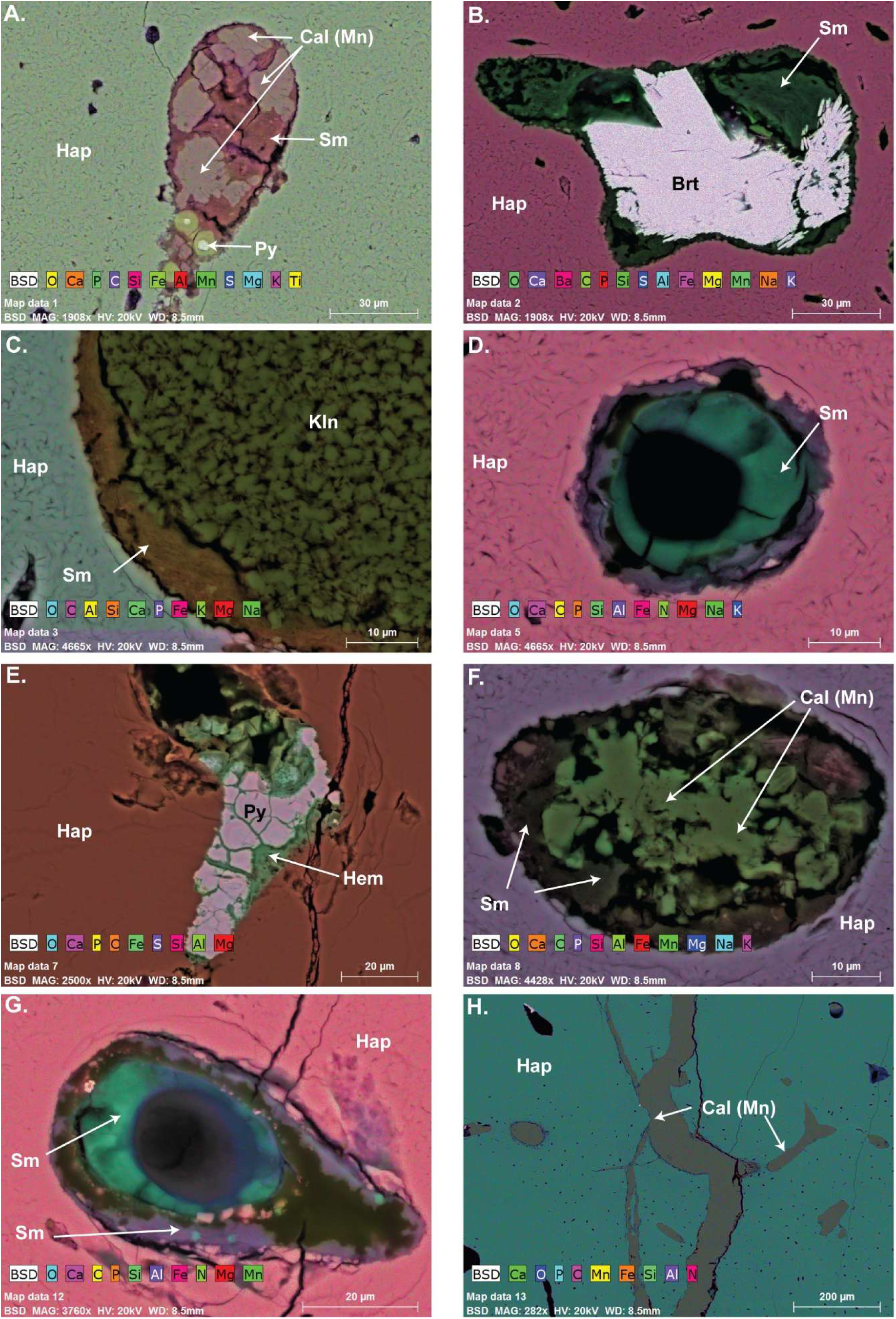
Variations of Haversian and Volkmann’s canal mineral infilling across the fibula cross-section as determined by EDS element distribution maps. A) The hydroxyapatite-based (Hap) bone matrix with an infilled Volkmann’s canal and manganese-dominant calcite (Cal (Mn)) and smectite-group minerals (Sm). B) Baryte (Brt) and smectite-group minerals (Sm) canal infilling within a hydroxyapatite-based matrix (Hap) C) kaolinite (Kln) and smectite D) smectite (Sm) infilling of a Haversian canal E) Hematite (Hem) and pyrite (Py) infilling of a crack region. F) Manganese dominant calcite (Cal (Mn)) and smectite-group minerals (Sm) infilling of a canal in the hydroxyapatite-based (Hap) matrix G) Smectite-group minerals (Sm) infilling of a canal H) Manganese dominant calcite (Cal (Mn)) mineralization in crack regions.

**Figure 3.**
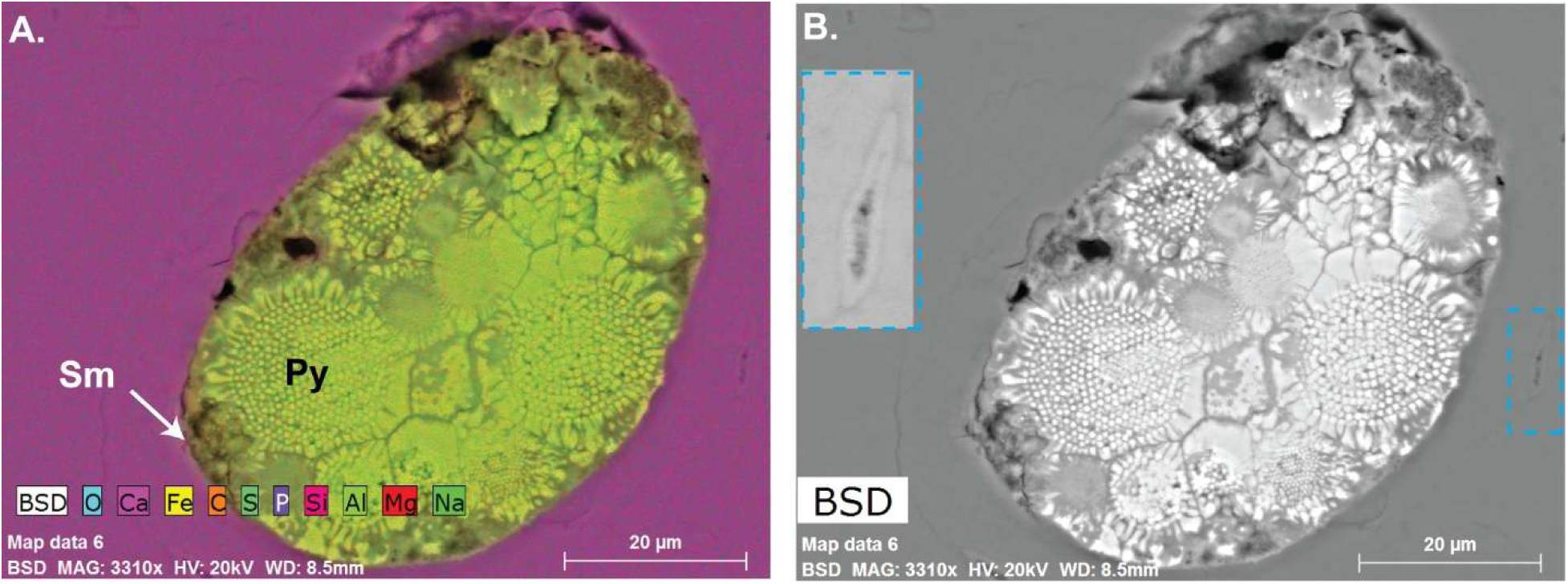
EDS element distribution maps show the pyritization and silicification in a Haversian canal in the cortex region of the Tyrannosaurid’s fibula. A) EDS map overlay of detected elements. B) Backscattered electron image reveals the filling of the haversian canal by framboidal pyrite (Py) structures embedded within smectite-group minerals (Sm). The inset in B) (blue dashed box) displays a mineralized lacuna in the cortex region containing calcium and phosphorus.

**Figure 4.**
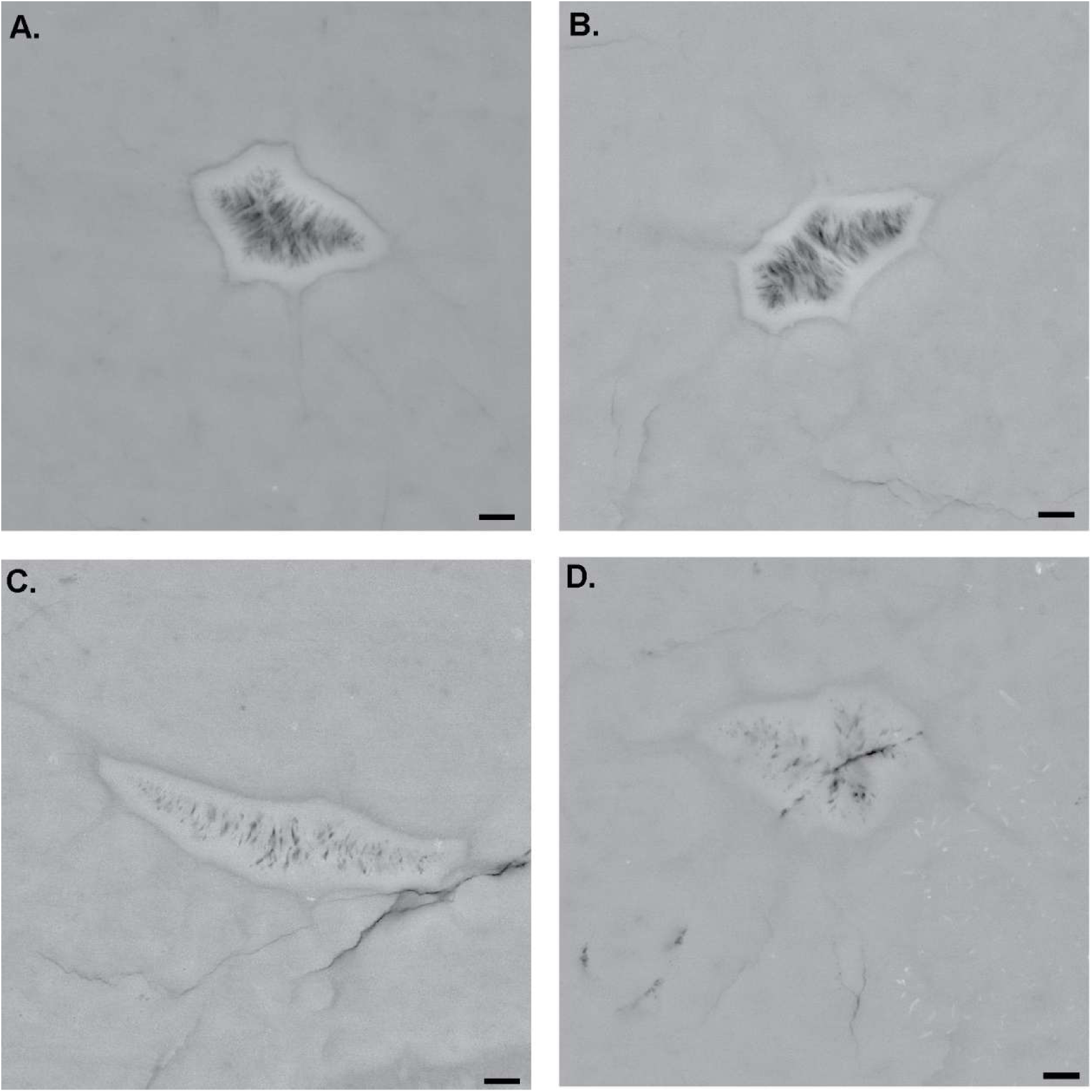
Backscattered electron images of lacunar mineralization in the cortex region which show CaP-based lamination and crystal structures. A) and B) Mineralized lacunae show a densely mineralized outer rim from which a network of needle-shaped crystals emanates into the center of the lacunae. EDS analyses (Figure 3B inset) reveal calcium and phosphorus as the main components. C) and D) display a higher degree of mineral infilling where the lacunae appear to be occluded. The crystal network is so dense and evolved that individual crystals cannot be distinguished from each other. Scale bar: 1µm

### 1.3 Evidence of Lacunar Mineralization

EDS results revealed that calcium and phosphorous mineralization in lacunae in the cortex (Fig 3B-Inset). Here mineral lamination was noted around the lacunar border with crystal structures protruding into the lacunar space (Fig. 4). Individual crystals were visible in some mineralized lacunae (Fig. 4A and B) in comparison to others (Fig. 4 C and D) where the mineral appears to occlude the entire lacunae, similar to what is seen during lacunar mineralization. Osteocyte lacunar mineralization has been termed “micropetrosis” ^82^, which has been visualized in modern (specimens younger than 100 years old)^83–89^ and fossilized^90^ specimens. Lacunar occlusions have also been visualized in human mandibles in high mineral-density regions, where neighbouring lacunae within the same osteon display different degrees of calcification^83^. Whether the mineralization is an active or passive process is still unknown. However, age^85,88^ and remodelling proximity^85^ have been noted as potential links to this process. Previous literature has found a significant increase in the number of mineralized lacunae from younger (< 39 years old) to older (> 80 years old) participants in a study population of the same number of human male and female specimens^85^. Inhibited or reduced remodelling, for example, as a result of cell death, may also impact lacunar mineralization. Mineralization factors produced by osteocytes will decrease upon their death, allowing for mineralization in the lacunar space^87,91^, which may initially occur in the pericellular region and grow into the remaining lacunar cavity^85^. Herein, mineralized osteocytes appeared to have minerals in both the lacunar and the canalicular space (Fig. 4). Studies in human bone have shown similarly that micropetrosis occurred in the lacunae and canaliculi, leading to the complete occlusion of the lacunae, creating a block in the LCN ^82,83,88^. Fossilized mammalian specimens up to 5 million years old have been shown to have mineral-filled lacunae, where remnants of cell structures, such as membranes and organelles, have been visualized previously in SEM and TEM analysis^90^. While cell structures were not identified from the 3D analysis in this study, further investigation using TEM could be implemented to verify the constituents inside the mineralized lacunae with higher resolution.

### 1.4 Collagen Fibrils Visualized Near the LCN

Across the transverse thin section of the fibula, multiple osteocyte lacunae were visible. In 2D backscatter SEM imaging of the medial and lateral regions of the fibula, lacunae have a dark appearance, where osteocytes were not visible in these spaces. Subsequent 3D FIB-SEM in the lateral region confirmed these lacunae as vacant (Fig. 5). The lacunar space was not infilled with epoxy resin during the embedding process, which created charging artifacts during 3D imaging. However, the lack of infiltration allowed for observations into the lacunae and visualization of key features, including pits that led to canaliculi and the disordered collagen fibril network that lined the lacunar space (Fig. 5 E, F). The lacunae are lined with mineralized collagen fibrils, forming a network in multiple directions (Fig. 5 E, F). This organization is similar to the loose and random arrangement of collagen fibres, termed osteocytic collagen, seen surrounding sectioned lacunae in human bone^92^. EDS map of this cross-section (Fig. S3) displayed elemental signatures of calcium, phosphorous, copper and gallium. The source of gallium is likely contamination from the FIB gallium-ion source, where the gallium can be redeposited onto the cross-section face during FIB milling^93^.

**Figure 5.**
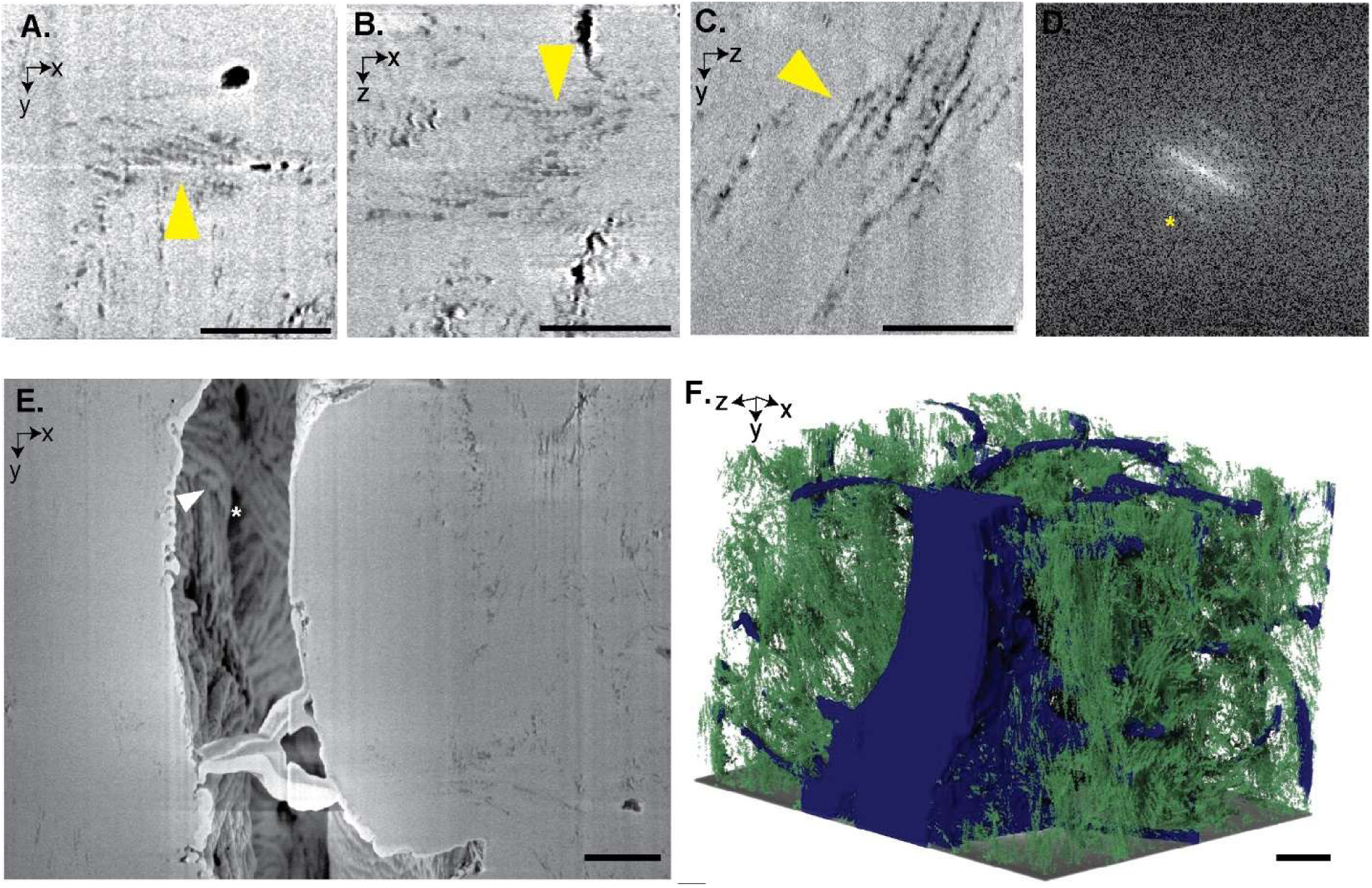
Orthogonal slices and 3D visualization of collagen and the lacunar-canalicular (LCN) space in the lateral region of the tyrannosaurid fibula acquired by FIB-SEM (#4 ROI-Fig S1). Collagen fibrils with distinct 67 nm D-banding periodicity were visualized in the A) *xy*, B) *xz* and *yz* orthogonal planes. The 67 nm periodicity of the collagen fibrils can beseen in the D) fast Fourier transform (FFT) image of C). E) An *xy* slice acquired with the secondary electron detector (SE2) of a lacunae where the pits (white asterisks) leading to the canaliculi and the CaP-covered collagen fibril network (white arrowhead) can be seen. Collagen in the plane ofthe lacunae is also seen in multidirections. F) 3D rendering of the LCN (blue) and collagen (green) where the collagen often appears to be co-aligned with the canaliculi. Scale bar: 1µm

In addition to surrounding the lacunae, collagen fibrils with ∼67 nm D-banding periodicity were seen in low mineralized regions across the bone matrix and in multiple imaging planes, i.e. the imaging plane collected in the FIB-SEM, and orthogonal reconstructed planes (Fig. 5. A-C). Fast Fourier Transforms (FFT) from the reconstructed *yz* image plane confirmed their ∼67 nm periodicity (Fig 5.). These results confirm the detection of intact individual collagen fibrils in their native mineral environment in this ∼71.5-million-year-old bone tissue specimen. Collagen fibrils surrounding and lining canaliculi were also found in alignment along their length (Fig. 6). This arrangement has been documented in FIB-SEM studies of modern specimens of human female femora^94^.

**Figure 6.**
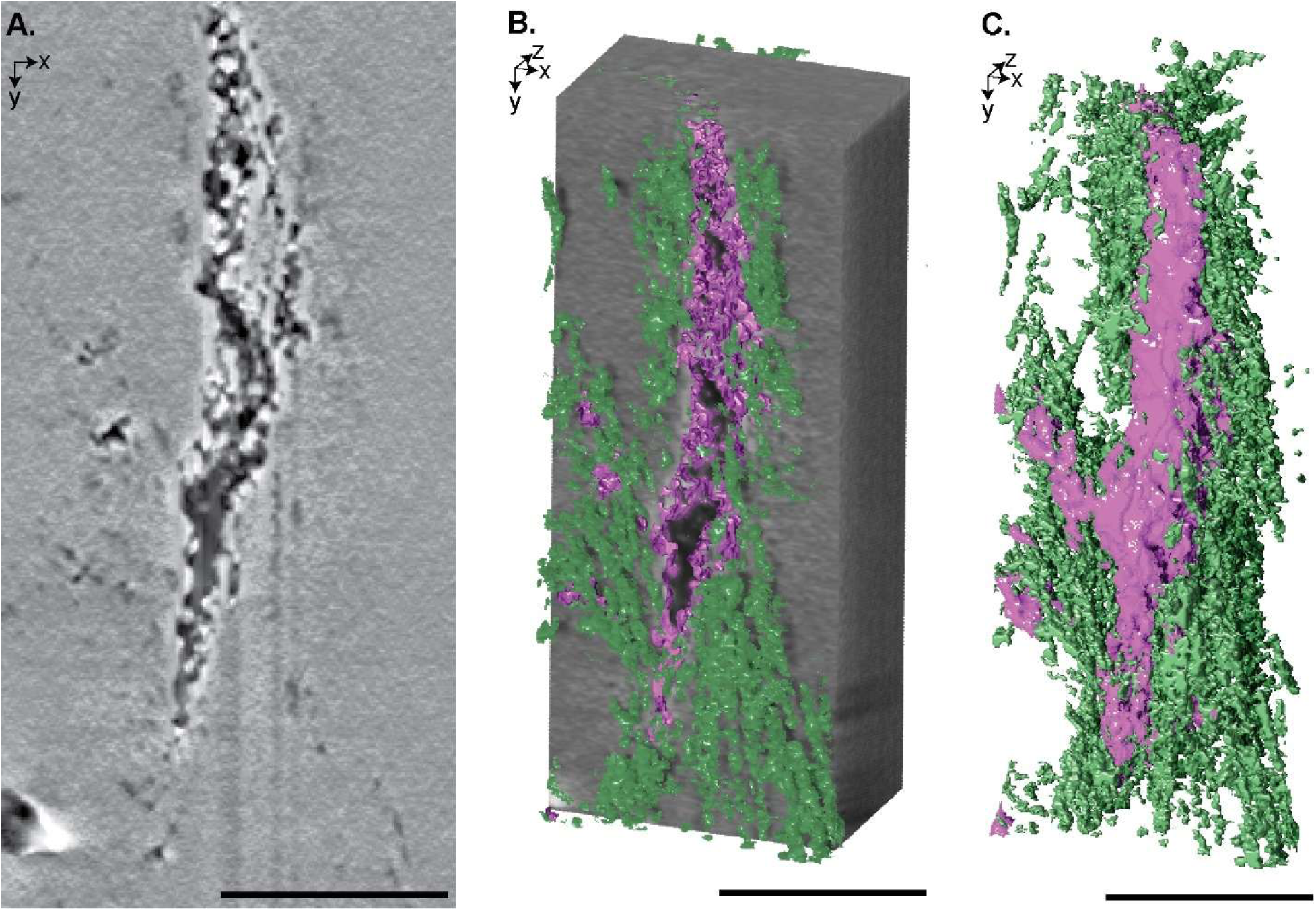
Cross-sectional and 3D visualization of collagen fibrils nearby and along the edge of a canaliculus from FIB-SEM data (#2 ROI-Fig S1). A) An *xy* slice of a canaliculus with circular features along the edge that correlate with collagen fibrils. B) 3D rendering of the collagen (green) and the outline of the canaliculus (pink) superimposed on the FIB-SEM volume. C) 3D rendering of just the collagen (green) and canaliculus (pink), showing their co-alignment. Scale bar: 1µm

Diagenetic processes change and degrade the organic matter in fossilized specimens^2,4^. However, there are records of soft tissue preservation in fossil bone^4^, including skin^13^, fibrous matrixes^10,15–17,21^, muscle^14^, DNA^95^, and other biomolecules^96^. In fossilized bone specimens, soft tissue, including fibrous matrix^9,10,15–17,97^, osteocytes^9,11,17–20,98^, and vessels^9,11,18,20^ have been visualized. Schweitzer et al. (2006)^9^ investigated demineralized bone samples across geological timescales from the Recent to the Cretaceous Period, showing a fibrous matrix that was previously associated with the mineral material in the bone. Demineralization has also revealed other soft, flexible tissue features, including osteocytes and the vasculature network within the bone tissue. For example, in sections of demineralized cortical bone from *Tyrannosaurus rex,* osteocytes are seen embedded within the fibrous matrix under light microscopy imaging^9^. Soft tissue preservation in fossilized specimens is frequently seen in environments containing fluvially derived sandstone ^9,10^. In the present study, the tyrannosaurid sample derives from the Horseshoe Canyon Formation in Alberta, Canada, which consists of interbedded sandstones, mudstones, and coaly layers, deposited by fluvial processes^99^. The preservation and composition of soft tissues in fossilized specimens are thought to be impacted by the tissue environment and secondary reactions, including cross-linkages to create decay-resistant organic compounds^4,17^. The fossil bone environment may also protect soft tissue features where the decay processes are slowed, thereby allowing for further reactions that can stabilize organic compounds^9,100,101^. Mineral-filled water infiltration from the local environment may pause organic decay in the fossils through external mineralization of the infiltrated minerals (ex., pyrite mineralization) ^4,12^. Changes from the potential original structure are seen in Paleolithic collagen, which varied biochemically from modern collagen with variations and loss in the amino acid composition^21^. Bertazzo et al. (2015)^16^ analyzed the collagen content in dinosaur specimens from the Cretaceous Period (∼71 million years ago), where FIB-SEM imaging revealed fibre fragments in low mineralized regions in the bone matrix. TEM analysis of these fragments revealed a fibrous structure with a ∼67 nm banding periodicity similar to collagen^16^. Time of flight secondary ion mass spectrometry (ToF-SIMS) analysis of regions containing these banded fibrous structures revealed amino acid structures, including glycine and proline^16^, confirming the presence of collagen fibrils in fossilized specimens dating to the Cretaceous ^16^. Other techniques that have been used for visualizing preserved collagen in fossilized specimens include immunofluorescence^10^, AFM^15^ and SFTIR^102^. In our investigation, the characteristic ∼67 nm periodicity of collagen fibrils was seen and accurately measured within the bone matrix (Fig 5. A-D). While mass spectrometry was not conducted in this study, the precise banding periodicity seen along fibrils displays a conservation of the quaternary structure of collagen^34,103,104^. Our findings support the preservation of collagen fibrils as an integral component of the bone matrix, while the disorganized collagen fibril arrangement around lacunae and alignment along canaliculi mirrors the patterns observed in modern species.

### 1.5 Organization of Collagen Fibril Bundles

SEM and FIB-SEM visualizations of the tyrannosaurid thin section showed evidence of collagen fibril bundles along the periosteal region. The lateral side of the fibula displayed several primary osteons with collagen fibril bundles, visible both proximally and distally to the Haversian canals in the fibrolamellar bone tissue (Fig. 7). Fibrolamellar bone is composed of a mix of bone tissue and is used to describe rapidly developing bone in modern and prehistoric animals^53,56–59,62,64,66,105–107^. Fibrolamellar bone tissue is characteristic of the fast-growing phase of juvenile mammals and dinosaurs, as this primary bone tissue is deposited rapidly and is eventually remodelled into more mature secondary lamellar bone^62,64,108^. The main components of fibrolamellar tissue include lamellar bone and non-lamellar bone (parallel-fibered bone and woven bone), where collagen fibrils in and outside bundles can be arranged longitudinally (parallel-fibered), randomly (woven bone) or in lamellae (lamellar bone)^62,64^.

**Figure 7.**
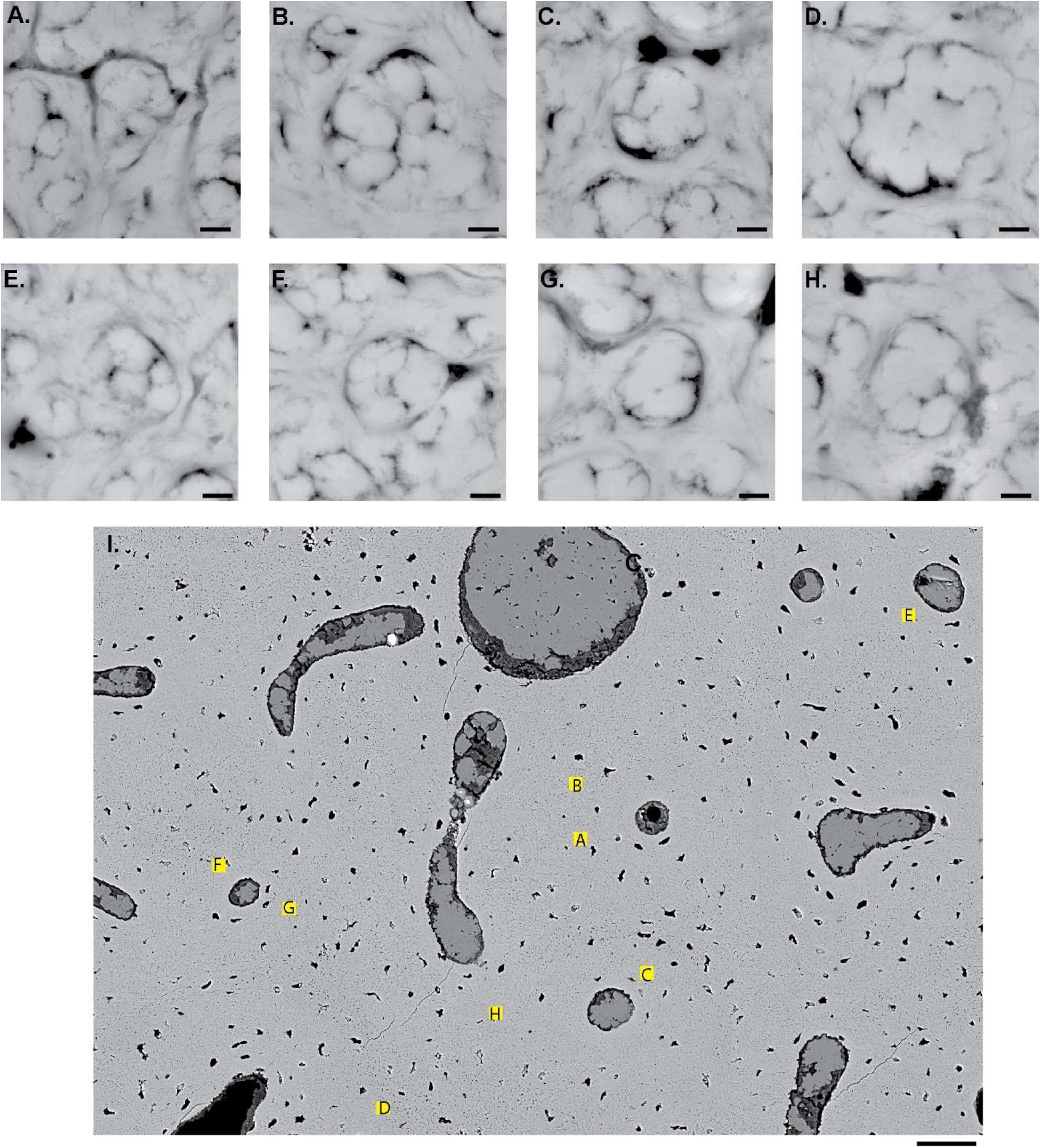
Backscattered electron images of collagen fibril bundles in the lateral region of tyrannosaurid fibula. A)-H) Collagen fibril bundles were delineated by either black features from either less mineralized collagen fibrils or canaliculi. I) Selected region of the large-area mosaic showing the locations of (A)-(H) where collagen fibril bundles were seen near and in between Haversian and Volkmann’s canals. Scale bars a)-h): 1 µm, i) 50 µm

Collagen fibril bundles were delineated by organic features or canaliculi (Fig. 7) and appeared to be 4.26 µm ± 1.06 in diameter, based on 2D analysis from the lateral side of the fibula. 3D FIB-SEM investigations visualized multiple arrangements of the collagen fibril bundles, including lamellar and non-lamellar arrangements (ex. parallel fibered arrangement) (Fig. 1). In 2D and 3D, canaliculi are present in and around bundles and did not appear to transect collagen fibril bundles but, in most cases, surround the collagen fibril bundles (Fig. 7). The fibril bundles were categorized based on the presentation of the low-mineralized collagen fibrils that could be visualized inside and around the bundles. Fibril bundles can range from 0.2 to 12 µm in diameter^104^. Investigations of human and minipig bone have revealed bundles between 2 and 3 µm in diameter^56,109^. Collagen bundles may either possess or lack an enclosing sheath^56,109,110^. Sheathed and unsheathed collagen fibril bundles can be visualized in the same specimen and even in the same bone type (ex. parallel fibered bone)^56^. Similar to bone tissue, in unmineralized turkey tendon tissue, a sheath or endototendon envelops collagen fibril bundles, and canaliculi appear to embed within the sheath structure^110^. While the mechanism and components for sheath development in bone are still relatively unknown, the development of the sheath regions may be linked to the bundle’s proximity to highly calcified regions seen in demineralized bone^56^. In this study, collagen fibril bundles in fibrolamellar bone do not appear to have sheath structures surrounding bundles but rather collagen fibril layers determining the bundle boundaries. The diagenesis processes may also impact the visualization of the sheath structures in the fossilized bone; specifically, changes to the mineral and organic content, including organic decay, mineral dissolution and recrystallization^1,2,5,14,21^. As such, it is difficult to conclusively state whether sheath structures around the collagen fibril bundles were previously present within the fossilized bone.

Collagen fibrils arranged in a parallel fibered arrangement are found in the middle of the bone matrix (Fig. 1D (Box 1) and Fig 9) and near a Volkmann’s canal (Fig. 1D (Box 2) and Fig. 8). In the parallel fibered tissue (Fig 8A and 9A), the collagen fibrils and canaliculi appeared to be aligned with the long axis of the fibula. Bundles are delineated by either collagen fibrils surrounding the bundle or canaliculi (Fig. 8B-C and 9 B-C), where the *xz* view displays the bundle cross-section (Fig. 8B and 9B). Orientation analysis on the resliced datasets from the *yz* plane displayed collagen fibrils and canaliculi that appeared to be longitudinally oriented (90 ° or -90°) (Fig 8 D-E and 9D-E). Non-longitudinally oriented collagen fibrils and canaliculi are also present, potentially due to the organic features delineating the bundles. These non-longitudinal features are also seen in small peaks or troughs in the dominant direction graph (Fig 8 E and 9. E). In the region near the Volkmann’s canal (Fig. 1D-Box 2), collagen fibrils within the bundle were difficult to visualize due to extensive mineralization; however, fibrils in the regions lining and surrounding the bundles appeared less mineralized (Fig. 9).

**Figure 8.**
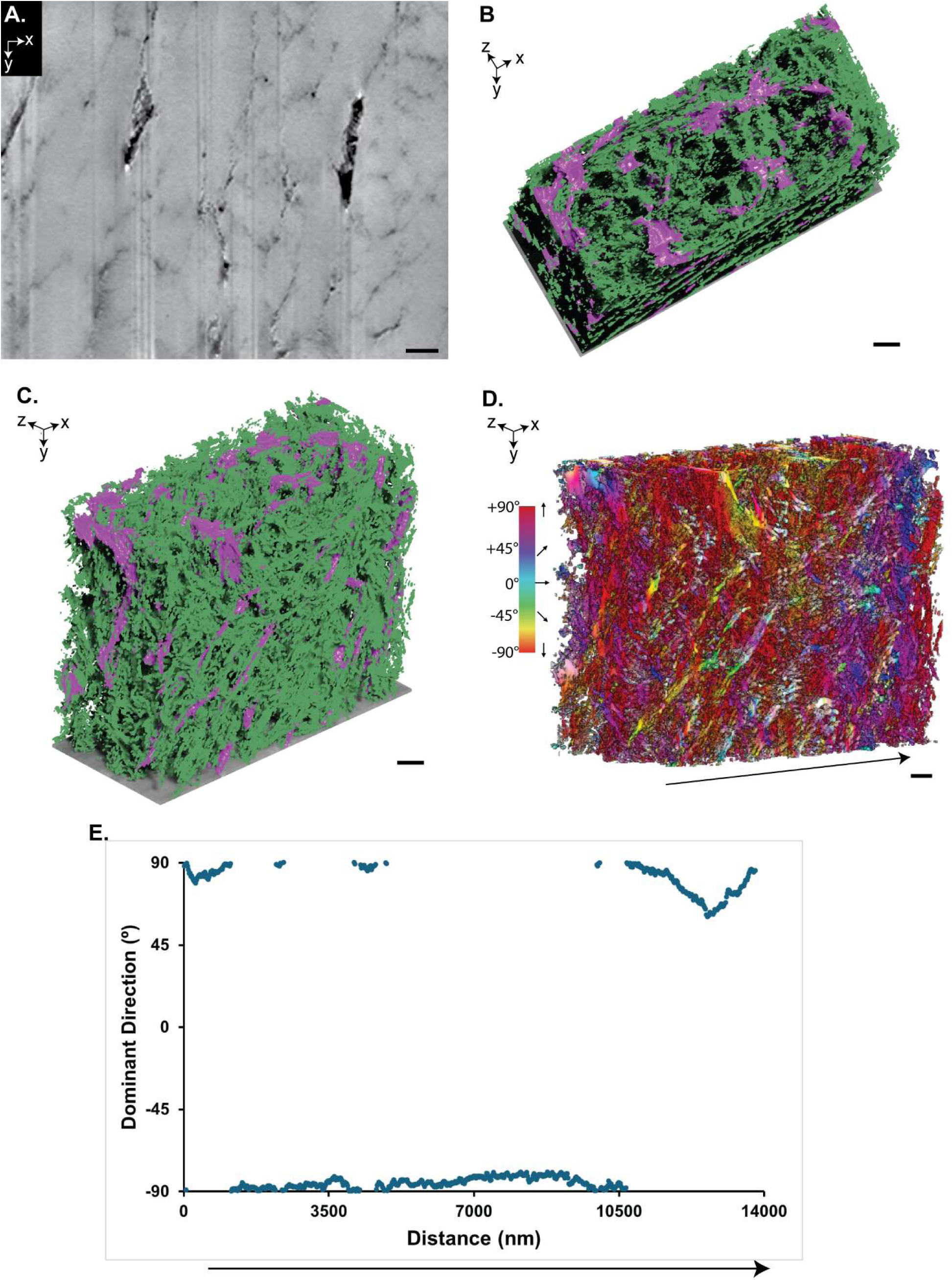
Parallel fibered bone array arrangement in the bone matrix (#1 ROI-Fig S1). A) An *xy* slice that runs longitudinally, parallel to the long axis of the bone, with mineral clusters visible, outlined by darker collagen fibrils or canaliculi. B) Top and C) side view of a 3D rendering of collagen (green) and canaliculi (pink). D) 3D orientation map of the individual collagen fibrils and canaliculi where the direction is indicated by a colour survey. Dominant direction analysis was conducted from the left to the right side of the volume (arrow) E) Dominant direction graph of the collagen fibrils and canaliculi orientation. Most fibrils and canaliculi are oriented either at 90^0^ or -90^0^. Scale bar: 1µm

**Figure 9.**
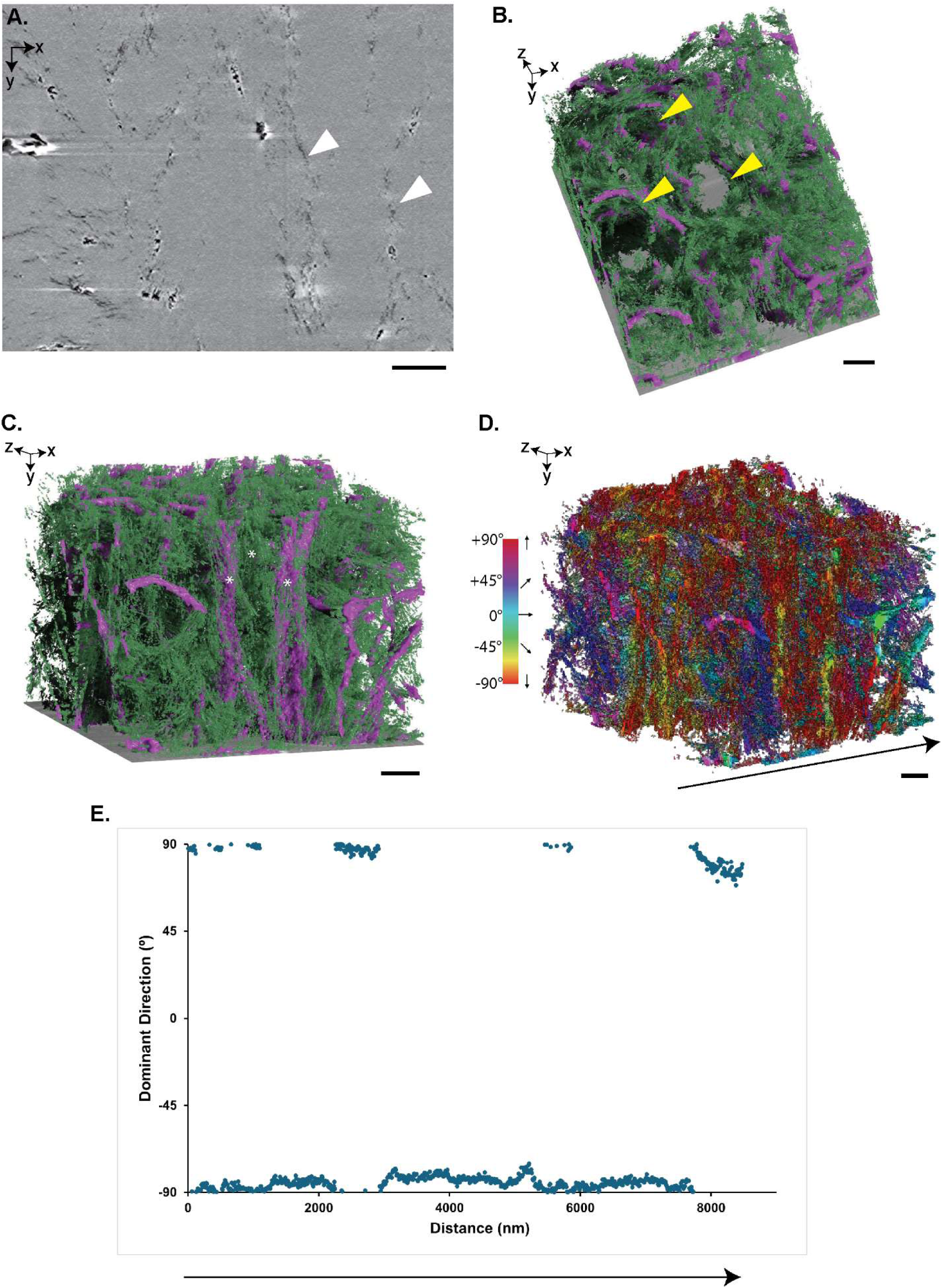
Parallel fibered bone array analysis near Volkmann’s canal (#2 ROI-Fig S1). A) An *xy* slice with collagen fibrils clear in darker regions (white arrowheads). B) Top and C) side view of 3D rendering of collagen (green) and canaliculi (pink). Collagen fibrils and canaliculi delineated the collagen fibril bundles (yellow arrowhead). Fibrils in the bundles were very highly mineralized and too obscure for segmentation. D) 3D orientation map of the collagen fibrils and canaliculi where the direction corresponds to the color survey. Fibrils and canaliculi were further analyzed from left to right (arrow) using dominant direction analysis and plotted. C) Dominant direction of fibrils and canaliculi, where the majority was oriented either at 90^0^ or -90^0^. Scale bar: 1µm

Near primary osteons, the collagen fibril organization resembled a lamellar bone configuration (Fig. 1D (Box 3) and Fig. 10). Collagen fibrils appeared to alternate between a longitudinal or transverse perspective (Fig. 10A). In this dataset, collagen fibrils inside and outside of the fibril bundles were visualized, as well as canaliculi (Fig. 10 B-C). Orientation analysis of individual collagen fibrils (Fig. 10 D) and dominant direction analysis (Fig. 10 E) from the resliced dataset from the *yz* plane, displayed fibrils that were oriented close to 85-90° or -85-90° on the left and right sides (Fig. 10 E-yellow region). However, some fibrils and canalicular structures are not oriented longitudinally in these regions. For example, there is a trough in the left yellow region that spans ∼750 nm (from slice 1000-1750, approximately); this change in the collagen and canaliculi direction could be due to a small layer of disorganized material in between the sub lamellae layers. The central area (Fig. 10-blue region) displays a gradual change in the direction of the collagen and canaliculi. This gradual change in fibril direction resembles the fanning collagen fibril array seen through FIB-SEM investigations in rat tibiae^111^. Plateau periods are also seen where the dominant direction of the fibrils and canaliculi remains the same (Fig 10. E). This smaller area resembles the unidirectional array of collagen fibrils seen within lamellar bone tissue^111^. While this provides insight into the organization of the collagen fibrils in fossilized specimens, a larger acquisition volume in this region could potentially display the alternating collagen fibril orientation to a more extensive degree than what was visualized here.

**Figure 10.**
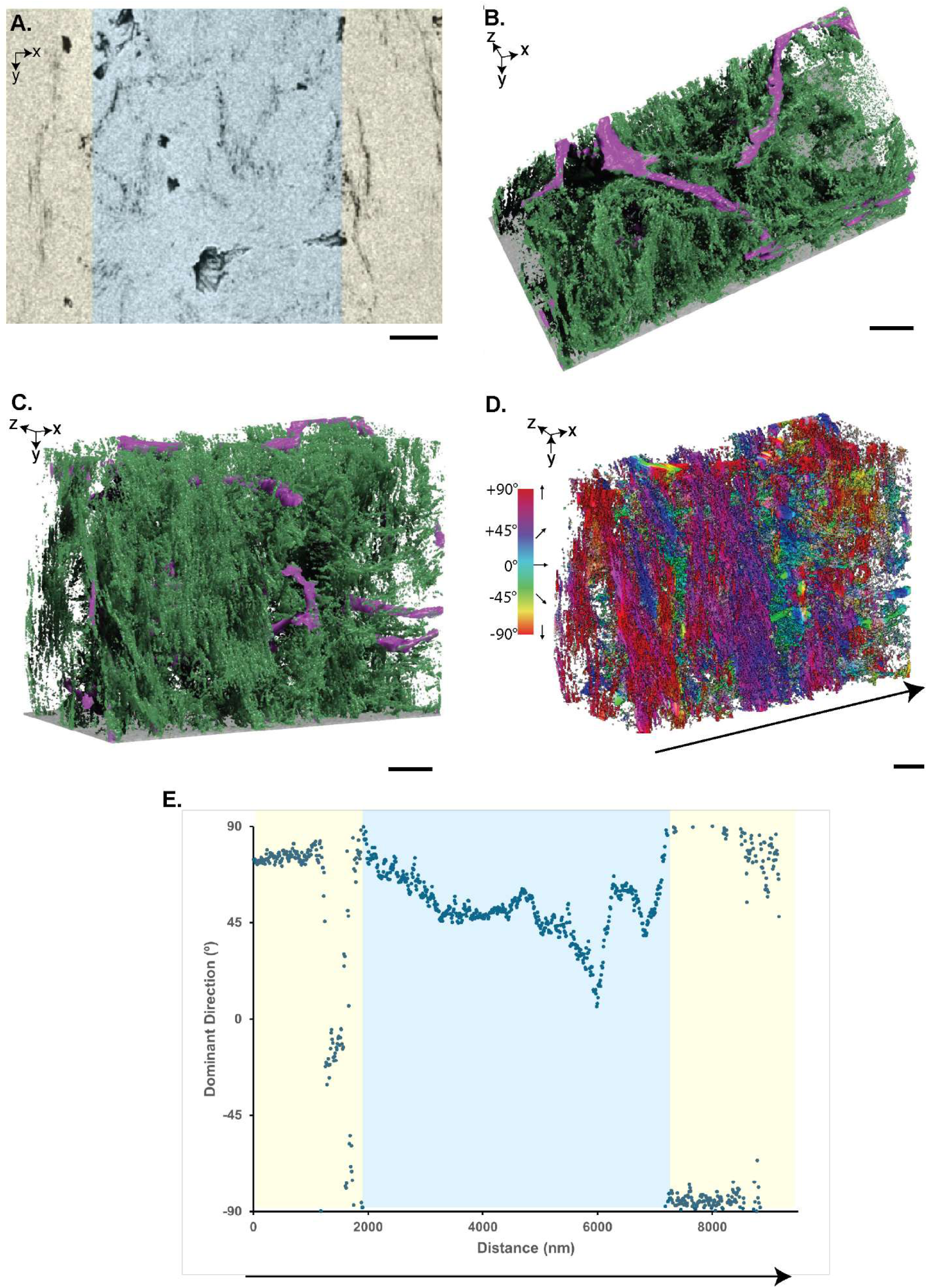
The lamellar fibered bone arrangement is visible near the Haversian canal (#3 ROI-Fig S1). A) An *xy* backscattered slice of collagen fibrils and canaliculi. Fibrils appear to be longitudinally oriented on the sides (yellow highlighted) and cross-sectionally oriented in the middle (blue) of the plane. B) Top and C) side view of 3D rendering of collagen (green) and canaliculi (pink). D) 3D orientation map of individual collagen fibrils and canaliculi with the direction indicated by the colour survey. Dominant direction analysis was performed on the FIB-SEM dataset from left to right (arrow) E) Dominant direction graph displaying fibrils and canaliculi as oriented near to or at 90^0^ or -90^0^ on the left and right sides of the dataset but having a “fanning” array in the middle displaying a gradual change in direction. Scale bar: 1µm

Previously, others investigated fibrolamellar bone units in demineralized minipig femoral bone using 3D FIB-SEM imaging, displaying a hypercalcified layer at the unit ends followed by parallel fibered and lamellar bone in the unit interior^56^. Fractured demineralized minipig bone also displayed sheathed collagen fibril bundles in the parallel-fibered bone with canaliculi surrounding bundles in some locations^56^. Similar ordered zones of parallel-fibered and woven bone with emerging lamellar bone have also been seen in subadult *Protoceratops andrewsi* using polarized light microscopy^58^. From our study, both parallel-fibered and lamellar bone are visible in the tyrannosaurid’s fibrolamellar bone tissue (Fig. 8-10). However, the hypercalcified layer seen in the demineralized minipig bone^56^ is not visible from either light microscopy, SEM or FIB-SEM imaging of the fossilized tyrannosaurid bone. 3D FIB-SEM analysis on multiple areas along the tyrannosaurid sample would be required to define its fibrolamellar unit structure conclusively. However, our results demonstrate the characteristic parallel-fibered and lamellar bone arrangement seen in fibrolamellar bone.

### 1.6 Mineral Clusters Revealed in Parallel Fibered Bone

In the parallel fibered bone investigated in the lateral region (Fig. 1D-Box 1), 1-3 µm long mineral clusters were found within the collagen bundles in the FIB-SEM dataset (Fig. 11). Mineral clusters were defined by surrounding organic features, including poorly mineralized collagen fibrils that defined the cluster’s boundaries. Using the watershed segmentation method^50,112^, 372 mineral clusters were segmented. The mean ferret diameter of the mineral clusters has a median of 1.91 µm and an interquartile range of 1.59 µm to 2.28 µm. The interquartile range of the aspect ratio mineral clusters is 0.33 to 0.53 for the aspect ratio with a median of 0.44, creating a 3D ellipsoidal mineral cluster shape (Fig. 12). These mineral clusters resemble mineral ellipsoids or “tesselles” recently discovered in human^37^ and vertebrate mammals^36,50^ bone. Some mineral clusters are larger than the average (Fig. 11C), which could be due to a lack of visible organic features at their periphery due to the high mineral density.

**Figure 11.**
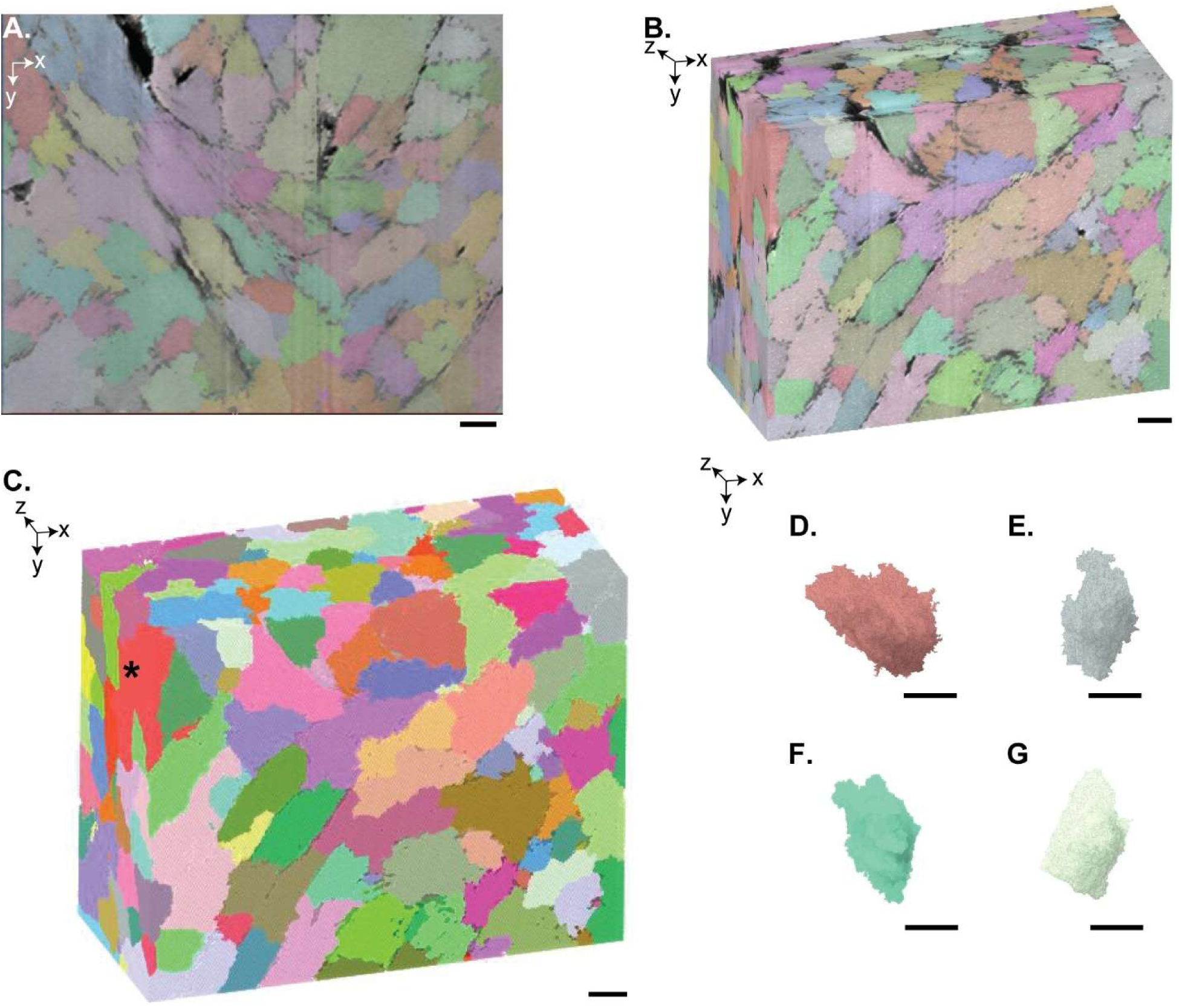
3D watershed segmentation and visualization of mineral clusters within parallel fibered bone (from Fig S1, ROI 1). A) SEM image in the *xy* plane of parallel fibered bone. Low or unmineralized collagen fibrils and canaliculi delineate mineral clusters which are approximately ∼1-3µm long. B) 3D overlay of watershed segmented mineral clusters and FIB-SEM dataset. C) 3D mineral cluster segmentation, where a few larger mineral aggregates are noted sporadically in the volume (black asterisk). D-G) Representative mineral clusters of approximately 1-3 µm mean ferret diameter and 0.3-0.6 aspect ratio displaying a 3D elliptical shape. Scale bar: 1µm, Colour scale: randomly assigned.

**Figure 12.**
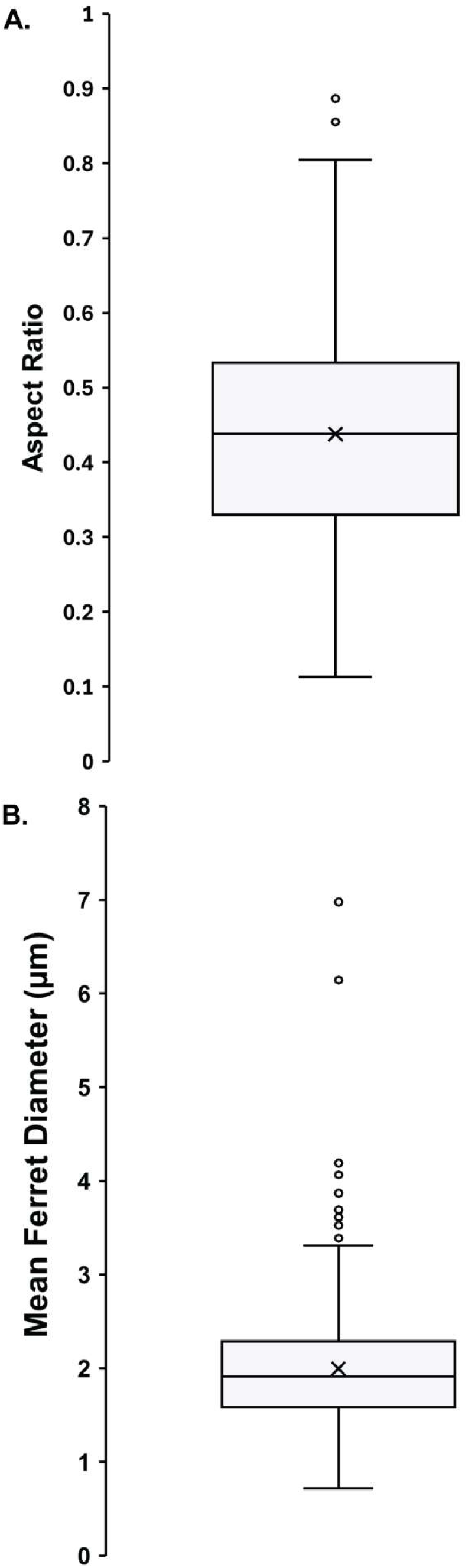
A) Box whisker plot of the aspect ratio of the segmented mineral clusters in the parallel fibered bone where the mean (0.44) is indicated by an x within the interquartile range (0.33-0.53) of the data and the median line (0.44) is shown. B) Box whisker plot of the mean ferret diameter of the segmented mineral clusters in the parallel fibered bone, where the mean (1.99 µm) and the interquartile range (1.59-2.28 µm) with a median of 1.91 µm is displayed (n=372).

In human and murine bone, mineral ellipsoids or “tesselles” are ellipsoid clusters of mineral, composed of carbonate-substituted hydroxyapatite mineral platelets^37,38,50^. Mineral ellipsoids are thought to develop from mineral foci within the collagen fibril network, where the mineral grows from the single point until it encounters its neighbouring mineral ellipsoid^36,113^. As the bone mineralizes, these ellipsoids become more defined and end with dimensions of approximately 750 nm in diameter and 1.5 µm in length^36,37,50^.The mineral clusters revealed by our investigation are slightly larger, on average, than the mineral ellipsoids seen in modern mammals. However, the overlap in size and shape of the mineral clusters suggests that they are homologous. To the authors’ knowledge, ours is the first account of the visualization of mineral ellipsoids within fossil specimens and within the parallel fibered bone pattern. This implies that nanoscale mineralization development in parallel and lamellar fibered bone samples may be similar. The difference between mineral ellipsoids from our investigation and modern investigations could stem from the differences in the bone pattern between parallel-fibered and lamellar bone tissue and from changes in the mineral during diagenesis. Hydroxyapatite can experience modifications to the crystal lattice structure through dissolution, substitution and recrystallization mechanisms, such as the ripening process^1,3,114,115^. The fluid flow of mineral-filled water from the environment can lead to the substitution and addition of external mineral ions (e.g., carbonate and fluoride) in the hydroxyapatite structure^1,3,115^, thereby influencing its recrystallization. The hydroxyapatite formed during diagenesis is larger compared to hydroxyapatite found in bones that have not undergone diagenesis^1,114,115^. Therefore, the morphology of the observed mineral clusters in the parallel fibered bone may be influenced by alterations in the mineral crystalline arrangement due to diagenesis. Despite these changes, however, it appears that during mineralization or recrystallization, mineral clusters appear to form a prolate ellipsoid shape akin to what has been seen in native modern-day bone.

### 1.7 Study Limitations

This investigation has provided details about the micro and nanostructure of the fibula of CMNFV 11315, including the permineralization process, mineralization of the lacunae, collagen fibril bundle arrangements, and mineral cluster arrangements. While we have accomplished enhanced visualization of these micro and nanoscale features, we acknowledge limitations in this study. A notable limitation is the number of FIB-SEM datasets acquired and analyzed. In this study, 4 FIB-SEM datasets are highlighted in the lateral region: 3 datasets reveal the collagen fibril pattern, and 1 dataset reveals the lacunar canalicular space. While these datasets provide insight into the collagen fibril network and arrangement in the fossilized bone, the limited number of datasets restricts broader conclusions that could be made about the fibrolamellar unit structure of the bone tissue. As such, more FIB-SEM datasets would need to be acquired and analysed to conclude the fibrolamellar unit structure. However, FIB-SEM remains a destructive analysis technique and therefore should be used with careful intention and restraint on such irreplaceable specimens.

## CONCLUSION

In this study, the micro and nanoscale osteohistological features of *Albertosaurus sarcophagus (*CMNFV 11315) were investigated, including Haversian canal and LCN permineralization, collagen fibril organization, and mineral cluster arrangements. Energy dispersive X-ray spectroscopy of this fossilized specimen showed evidence of diagenetic processes and permineralization, where external minerals infiltrated the fossil through fluid flow processes and settled to create secondary mineral structures, including smectite-minerals, framboidal pyrite and baryte crystals. Using FIB-SEM nanotomography, we resolved the preserved collagen fibril network in low mineralized regions, around cell processes and within the cell lacunae of this 70+ million-year-old specimen. In collagen fibril segments, characteristic ∼67 nm D-banding periodicity was visualized in the imaging and reconstructed orthogonal planes in the bone matrix environment. 3D FIB-SEM of fossilized bone tissue revealed different material patterns within the fibrolamellar tissue including lamellar and parallel fibered bone. Ellipsoidal mineral clusters were visualized in parallel fibered bone where clusters appeared to be in alignment with the collagen fibrils. Using 3D FIB-SEM, this investigation displays the first record, to the authors’ knowledge, of an extensive 3D collagen fibril network, and prolate ellipsoidal mineral clusters in a 71.5-million-year-old fossilized bone specimen. FIB-SEM nanotomography has proved itself a valuable tool for advancing our understanding of biomineralization and fossilization processes.

## MATERIALS AND METHODS

### Thin Section Details

Mallon et al. (2020)^61^ reported on the partial skeleton of a late juvenile– early subadult *Albertosaurus sarcophagus* (Canadian Museum of Nature [CMN] catalogue number FV 11315), which they subjected to osteohistological analysis for ontogenetic age determination. The authors estimated a minimum age at time of death of 2 years, based on the number of growth annuli, although they did not retrocalculate earlier annuli possibly obliterated by bone remodeling. The thin section they prepared is from the distal third section of the left fibula. This bone was chosen since it was broken post-mortem and originally bore little weight (hence, was presumably not so prone to related secondary remodeling^61^). The thin section was created by sectioning a 5 mm thick sample from the bone using a Buehler Isomet 1000 Precision Saw; the sample was mounted using Palouse Petropoxy 154 with hardener on a glass slide with epoxy and polished with an increasing polishing series. We repurposed this thin section for the present study. Fossils were collected, sampled and imaged with the appropriate permissions.

### Light Microscopy and Scanning Electron Microscopy

Large area transmitted light microscopy image mosaics were acquired using a Zeiss AXIO Zoom.V16 light microscope. The mosaics were acquired with ZEN Pro imaging software using a Plan Apo Z 1.0/0.25 objective (FWD 60 mm) at a resolution of 410 nm/pixel in plane-polarized (PPL) and cross-polarized transmitted light (XPL). The mosaics that were obtained consisted of 522 individual tiles. Large-area scanning electron microscope (SEM) image mosaics were acquired from the thin section CMNFV 13115 by using a Zeiss Gemini 450 field-emission (FE) scanning electron microscope and the software Zeiss Atlas 5. The light microscopy mosaics were imported into the Atlas 5 correlative workspace and aligned with the sample in the microscope. A large-area SEM overview mosaic was acquired at an acceleration voltage of 20 kV by using both the backscatter electron detector (BSD4) and the secondary electron (SE2) detector, a working distance of 10 mm, a 3.2 nA beam current, a 3.0 µs dwell time, and a resolution of 65 nm/pixel. The resulting mosaics comprise 834 image tiles each, with each tile consisting of 10240 x 10240 pixels (666.7 x 666.7 µm) and a total pixel count of 87.5 gigapixels. These large area light and electron microscopy mosaics were previously published in the prior microscale characterization of the CMNFV 11315 specimen by Mallon et al (2020)^61^ and segments of them are reproduced here with permission to enable locating regions of interest for elemental and 3D analysis, described below. High-resolution image mosaics and images were acquired at 10 nm/pixel from selected areas of interest on the fibula thin section. The mosaic was acquired at a pixel resolution of 10 nm, a dwell time of 5 µs, and a line averaging of 1. Once the image mosaics were acquired, stitched, and an image correction was performed, the entire Atlas 5 data set was exported to an autonomous series of files called the Browser-Based Viewer (BBV), which allows anyone using a PC, tablet, or cell phone to view the complete data set at full resolution in a web browser. The computer mouse is used to zoom in and out as well as to navigate through the large-area image mosaic. The data can be accessed at: https://www.petapixelproject.com/mosaics/museumofnature/CMNFV-11315/index.html

### Energy-Dispersive X-ray Spectroscopy

Energy-dispersive X-ray spectroscopy (EDS) was conducted to identify minerals in key regions of the CMNFV 11315 thin section. EDS imaging was performed on a Zeiss EVO MA 15 tungsten-filament SEM equipped with two Bruker XFlash 6/30 EDS detectors controlled using the Esprit 2.2 software. An acceleration voltage of 20 kV and a probe current of 3.7 nA were used for the acquisition of EDS element-distribution maps and point analyses. EDS analysis on one cross-sectional face of one Atlas 3D nanotomography run was performed on a Zeiss Crossbeam 540 FIB-SEM at 20kV and 54° stage tilt using the Oxford Instruments Aztec EDS software. All element-distribution maps and point analyses were exported from the Bruker Esprit and the Oxford Aztec software, arranged into figure plates using the software CorelDraw 24 and Adobe Photoshop, exported as PDF or JPG files, and linked with their respective location of acquisition in the Atlas 5 browser-Based Viewer dataset.

### FIB-SEM Nanotomography

FIB-SEM serial sectioning was performed using the Zeiss Atlas3D: Nanotomography software (Fibics, Ottawa, Canada) on a Zeiss Crossbeam 350 FIB-SEM. FIB-SEM nanotomography regions of interest (Fig 1D, red boxes) were in the lateral region where the red arrow indicates the direction of the imaging progression, or sequential image stack. Resolution and imaging details for each dataset are detailed in Table S1, while videos are also attached to the Atlas 5 browser-Based Viewer dataset.

The A3D nanotomography preparation included depositing a protective tungsten or platinum layer on the region of interest, then milling 3D tracking and autotune fiducial marks to perform automated tracking and autofocus. Milled fiducial marks were highlighted with a deposited carbon layer, and a final carbon protective layer was deposited onto the entire preparation pad. High FIB probe currents (ex. 30 kV; 30 nA and 3 nA) created stepped trenches in the specimen to expose the 3D imaging surface. Automatic serial sectioning and imaging was then performed using the appropriate imaging and milling probe (Table S1). The imaging signal from the secondary electron detector (SE2) and the energy-selective backscattered electron (EsB) detector were collected. Both the SE2 and EsB datasets were processed to enhance contrast in FIJI (ImageJ v1.53t) (NIH)^116^ using the “Enhance Local Contrast (CLAHE)” macro for further analysis. Applied CLAHE parameters included a block size of 200, a bin of 256 and a slope of 3. Fast Fourier transform analysis was conducted in FIJI using the FFT plugin.

### Image Alignment and Segmentation

FIB-SEM datasets processed using CLAHE were imported into Dragonfly (Version 2022.2) (Object Research Systems, Montreal, Canada). Slices were registered using the “Sum of Squared Differences (SSD)” and Mutual Information algorithms to align the datasets. The SE2 signal from the dataset was used for image analysis except for one dataset specified in Table S1, where the EsB signal was used instead due to compounded imaging artifacts (i.e. curtaining and out-of-focus images) present in the SE2 dataset. Vertical de-striping was used to remove curtaining artifacts seen in some of the SE2 FIB-SEM datasets. Gaussian smoothing aided in denoising the images to improve feature visualization. Slope map was also implemented for the second dataset to improve the imaging and shading compensation was also applied to compensate for shadowing that occurred during FIB-SEM imaging. A summary of the image processing steps pertaining to the relevant runs are detailed in Table S2.

Lacunocanalicular space, canaliculi and/or collagen fibrils were segmented in FIB-SEM datasets using a U-Net model. The model was trained in the segmentation wizard module of Dragonfly to identify the organic features within bone. Approximately 10-15 frames were manually segmented using the ROI painter tool to identify the features of interest. These frames were used to train the U-Net model to segment these features within the FIB-SEM datasets. The U-Net model parameters included a patch size of 64, batch size of 128, stride ratio of 0.25, 100 epochs for training and a learning rate of 1. The model was considered trained and appropriate for use if it had achieved an ORS DICE loss score of 0.9 or greater. The trained model was applied to the entire dataset to segment the collagen and canaliculi. U-Net models were trained to identify features for each dataset.

A watershed segmentation was used to identify mineral clusters in the bone. Sole thresholding segmentation did not segment mineral clusters as they overlapped, preventing cluster separation. As such, we used a previously reported method^50,112^ to segment the mineral clusters to identify their volume and shape distribution. An inverted distance map was created using the collagen and canaliculi segmentation. The centers of the mineral clusters were segmented by thresholding on the inverted distance map and applying the “open” tool (kernel size: 11) on the ROI. This segmentation created the “seeds” for the watershed segmentation. A multi-ROI of the individual seeds was created by separating the segmentation results based on the 6-connected components. The inverted distance map and the multi-ROI of the seeds were used as the inputs for the watershed transform. With the transformation, each seed for the mineral cluster was enlarged until it reached the boundary of another mineral cluster. The collagen and canaliculi segmentations were subtracted from the mineral cluster ROI using Boolean operations to avoid overlapping between the features during volumetric analysis.

### Dominant Direction and Orientation Analysis

The “OrientationJ” plugin^116–119^ was used within FIJI for orientation and directionality analysis. The datasets were first resliced along the *yz* plane to view the collagen fibrils within the same longitudinal view. The OrientationJ Dominant Direction tool^118^ was used to identify the dominant direction of the collagen fibrils within the same slice. The OrientationJ Analysis tool^117^ was used with a local window σ of 10 pixels, a gaussian gradient and an HSB colour survey where the hue represents the orientation of the fibrils.

## ACKNOWLEDGEMENTS

The Canadian Museum of Nature is gratefully acknowledged for providing CMNFV 11315 for 2D and 3D nanoscale analysis. Financial support was received from the Human Frontier Science Program Research Grant, the Natural Sciences and Engineering Research Council of Canada, and the Canada Research Chairs Program. Fibics Incorporated is acknowledged for 3D FIB-SEM nanotomography data acquisition and optimization.

## ASSOCIATED CONTENT

### Availability of data and material

Mosaics, elemental maps and videos of FIB-SEM acquisition can be found: https://www.petapixelproject.com/mosaics/museumofnature/CMNFV-11315/index.html Raw data is available upon request

### Supporting information

Supporting figures include high-resolution imaging of permineralization events, element distribution maps showing pyritization and silification and elemental maps of a lacuna cross-section. Supporting tables detail imaging parameters for FIB-SEM acquisitions and image processing steps of FIB-SEM data. Video captions of Video S1-5 are detailed in the supporting information file.

## AUTHOR INFORMATION

### Authors’ Contributions

Conceptualization: AW, DS, JM, NB, KG Methodology: AW, DS, NB, JM, KG, Data acquisition and formal analysis: AW and DS Supervision and financial support: KG, JM, NB, MP Writing: AW, DS, KG Review and Editing: ALL

### Funding Sources

Human Frontier Science Program Research Grant RGP0023/2021 to KG

NSERC Discovery Grant to KG (RGPIN-2020-05722) and NB

NSERC-CGSD and CGSM to AW

Canada Research Chairs Program to KG

## REFERENCES

1. Hedges, R. E. M. Bone diagenesis: an overview of processes. Archaeometry 44, 319–328 (2002).

2. Kendall, C., Eriksen, A. M. H., Kontopoulos, I., Collins, M. J. & Turner-Walker, G. Diagenesis of archaeological bone and tooth. *Palaeogeogr., Palaeoclim.*, Palaeoecol. 491, 21–37 (2018).

3. Sousa, D. V. de, Eltink, E., Oliveira, R. A. P., Félix, J. F. & Guimarães, L. de M. Diagenetic processes in Quaternary fossil bones from tropical limestone caves. Sci Rep-uk 10, 21425 (2020).

4. Briggs, D. E. G. The Role Of Decay and Mineralization in the Preservation of Soft-Bodied Fossils. Annu. Rev. Earth Planet. Sci. 31, 275–301 (2003).

5. Keenan, S. W. & Engel, A. S. Early diagenesis and recrystallization of bone. Geochim. Cosmochim. Acta 196, 209–223 (2017).

6. Pflug, H. D. Organic Geo- and Cosmochemistry. Top. Curr. Chem. 1–55 (2005) doi:10.1007/bfb0018077.

7. Chinsamy-Turan. The Microstructure of Dinosaur Bone : Deciphering Biology with Fine-Scale Techniques. (The Johns Hopkins University Press, 2005).

8. Farrell, Ú. C. Pyritization of Soft Tissues in the Fossil Record: An Overview. Paléontol. Soc. Pap. 20, 35–58 (2014).

9. Schweitzer, M. H., Wittmeyer, J. L. & Horner, J. R. Soft tissue and cellular preservation in vertebrate skeletal elements from the Cretaceous to the present. Proc. R. Soc. B: Biol. Sci. 274, 183–197 (2007).

10. Voegele, K. K. et al. Soft Tissue and Biomolecular Preservation in Vertebrate Fossils from Glauconitic, Shallow Marine Sediments of the Hornerstown Formation, Edelman Fossil Park, New Jersey. Biology 11, 1161 (2022).

11. Schweitzer, M. H., Wittmeyer, J. L., Horner, J. R. & Toporski, J. K. Soft-Tissue Vessels and Cellular Preservation in Tyrannosaurus rex. Science 307, 1952–1955 (2005).

12. Allison, P. A. The role of anoxia in the decay and mineralization of proteinaceous macro-fossils. Paleobiology 14, 139–154 (1988).

13. Manning, P. L. et al. Mineralized soft-tissue structure and chemistry in a mummified hadrosaur from the Hell Creek Formation, North Dakota (USA). Proc. R. Soc. B: Biol. Sci. 276, 3429–3437 (2009).

14. McNamara, M. et al. Organic preservation of fossil musculature with ultracellular detail. Proc. R. Soc. B: Biol. Sci. 277, 423–427 (2009).

15. Avci, R. et al. Preservation of Bone Collagen from the Late Cretaceous Period Studied by Immunological Techniques and Atomic Force Microscopy. Langmuir 21, 3584–3590 (2005).

16. Bertazzo, S. et al. Fibres and cellular structures preserved in 75-million–year-old dinosaur specimens. Nat. Commun. 6, 7352 (2015).

17. Senter, P. Cells and soft tissues in fossil bone: A review of preservation mechanisms, with corrections of misconceptions. *Palaeontol*. Electron. (2022) doi:10.26879/1248.

18. Pawlicki, R., Korbel, A. & Kubiak, H. Cells, Collagen Fibrils and Vessels in Dinosaur Bone. Nature 211, 655–657 (1966).

19. Pawlicki, R. Morphological differentiation of the fossil dinosaur bone cells. Cells Tissues Organs 100, 411–418 (1978).

20. Pawlicki, R. & Nowogrodzka-Zagórska, M. Blood vessels and red blood cells preserved in dinosaur bones. Ann. Anat. - Anat. Anz. 180, 73–77 (1998).

21. Tuross, N. Alterations in fossil collagen. Archaeometry 44, 427–434 (2002).

22. Kizilyaprak, C., Stierhof, Y.-D. & Humbel, B. M. Volume microscopy in biology: FIB-SEM tomography. Tissue Cell 57, 123–128 (2019).

23. Raguin, E., Rechav, K., Shahar, R. & Weiner, S. Focused ion beam-SEM 3D analysis of mineralized osteonal bone: lamellae and cement sheath structures. Acta Biomater. 121, 497–513 (2021).

24. Schneider, P., Meier, M., Wepf, R. & Müller, R. Serial FIB/SEM imaging for quantitative 3D assessment of the osteocyte lacuno-canalicular network. Bone 49, 304–11 (2011).

25. Weiner, S. & Wagner, H. D. The Material Bone: Structure-Mechanical Function Relations. Annual Review of Material Science 28, 271–298 (1998).

26. Fratzl, P. & Weinkamer, R. Nature’s hierarchical materials. Progr. Mater. Sci. 52, 1263– 1334 (2007).

27. Reznikov, N., Shahar, R. & Weiner, S. Bone hierarchical structure in three dimensions. Acta Biomater 10, 3815–3826 (2014).

28. Shah, F. A., Ruscsák, K. & Palmquist, A. 50 years of scanning electron microscopy of bone – A comprehensive overview of the important discoveries made and insights gained into bone material properties in health, disease, and taphonomy. Bone Res 7, 15 (2019).

29. Wess, T. J. Collagen, Structure and Mechanics. 49–80 (2008) doi:10.1007/978-0-387-73906-9_3.

30. Sorokina, L. V., Shahbazian-Yassar, R. & Shokuhfar, T. Collagen biomineralization: pathways, mechanisms, and thermodynamics. Emergent Mater 4, 1205–1224 (2021).

31. Kadler, K. E., Holmes, D. F., Trotter, J. A. & Chapman, J. A. Collagen fibril formation. Biochem J 316, 1–11 (1996).

32. Fratzl, P., Gupta, H. S., Paschalis, E. P. & Roschger, P. Structure and mechanical quality of the collagen– mineral nano-composite in bone. J Mater Chem 14, 2115–2123 (2004).

33. Cui, F.-Z., Li, Y. & Ge, J. Self-assembly of mineralized collagen composites. Mater Sci Eng R Reports 57, 1–27 (2007).

34. Orgel, J. P. R. O., Irving, T. C., Miller, A. & Wess, T. J. Microfibrillar structure of type I collagen in situ. Proc National Acad Sci 103, 9001–9005 (2006).

35. Hodge, A. J. & Petruska, J. A. Recent studies with the electron microscope on ordered aggregates of the tropocollagen macromolecule. in Aspects of protein structure (ed. Ramachandran, G. N.) 289–300 (Academic Press, New York, NY, 1963).

36. McKee, M. D., Buss, D. J. & Reznikov, N. Mineral tessellation in bone and the stenciling principle for extracellular matrix mineralization. J Struct Biol 214, 107823 (2022).

37. Binkley, D. M., Deering, J., Yuan, H., Gourrier, A. & Grandfield, K. Ellipsoidal mesoscale mineralization pattern in human cortical bone revealed in 3D by plasma focused ion beam serial sectioning. J Struct Biol 212, 107615 (2020).

38. Buss, D. J., Kröger, R., McKee, M. D. & Reznikov, N. Hierarchical organization of bone in three dimensions: A twist of twists. J Struct Biol X 6, 100057 (2022).

39. Clarke, B. Normal bone anatomy and physiology. Clin J Am Soc Nephro 3, S131–S139 (2008).

40. Lin, X., Patil, S., Gao, Y.-G. & Qian, A. The bone extracellular matrix in bone formation and regeneration. Front. Pharmacol. 11, 757 (2020).

41. Pritchard, J. J. The Biochemistry and Physiology of Bone. 179–212 (1956) doi:10.1016/b978-1-4832-3286-7.50011-7.

42. Robling, A. G. & Bonewald, L. F. The osteocyte: New insights. Annu Rev Physiol 82, 485– 506 (2020).

43. Atkins, G. J. & Findlay, D. M. Osteocyte regulation of bone mineral: a little give and take. Osteoporosis Int 23, 2067–2079 (2012).

44. Oers, R. F. M. van, Wang, H. & Bacabac, R. G. Osteocyte Shape and Mechanical Loading. Curr. Osteoporos. Rep. 13, 61–66 (2015).

45. Qing, H. & Bonewald, L. F. Osteocyte Remodeling of the Perilacunar and Pericanalicular Matrix. Int J Oral Sci 1, 59–65 (2009).

46. Bonewald, L. F. The amazing osteocyte. JBMR 26, 229–238 (2011).

47. Hemmatian, H., Bakker, A. D., Klein-Nulend, J. & Lenthe, G. H. van. Aging, Osteocytes, and Mechanotransduction. Curr. Osteoporos. Rep. 15, 401–411 (2017).

48. Tang, T. et al. A 3D Network of Nanochannels for Possible Ion and Molecule Transit in Mineralizing Bone and Cartilage. Adv Nanobiomed Res 2, 2100162 (2022).

49. Weiner, S., Raguin, E. & Shahar, R. High resolution 3D structures of mineralized tissues in health and disease. Nat Rev Endocrinol 17, 307–316 (2021).

50. Buss, D. J., Reznikov, N. & McKee, M. D. Crossfibrillar mineral tessellation in normal and Hyp mouse bone as revealed by 3D FIB-SEM microscopy. J. Struct. Biol. 212, 107603 (2020).

51. Sarathchandra, P., Pope, F. M., Kayser, M. V. & Ali, S. Y. A light and electron microscopic study of osteogenesis imperfecta bone samples, with reference to collagen chemistry and clinical phenotype. J. Pathol. 192, 385–395 (2000).

52. Giraud-Guille, M. M. Twisted plywood architecture of collagen fibrils in human compact bone osteons. Calcif. Tissue Int. 42, 167–180–167–180 (1988).

53. Whitney, M. R., Otoo, B. K. A., Angielczyk, K. D. & Pierce, S. E. Fossil bone histology reveals ancient origins for rapid juvenile growth in tetrapods. *Commun*. Biol. 5, 1280 (2022).

54. Windholz, G. J. et al. Osteohistology of Uberabatitan ribeiroi (Dinosauria, Sauropoda) provides insight into the life history of titanosaurs. Hist. Biol. **ahead-of-print**, 1–11 (2023).

55. Organ, C. L. & Adams, J. The histology of ossified tendon in dinosaurs. J. Vertebr. Paléontol. 25, 602–613 (2005).

56. Magal, R. A., Reznikov, N., Shahar, R. & Weiner, S. Three-dimensional structure of minipig fibrolamellar bone: adaptation to axial loading. J Struct Biol 186, 253–64 (2014).

57. Barrera, J. W., Cabec, A. L. & Barak, M. M. The orthotropic elastic properties of fibrolamellar bone tissue in juvenile white-tailed deer femora. J Anat 229, 568–576 (2016).

58. Fostowicz-Frelik, L. & Słowiak, J. Bone histology of Protoceratops andrewsi from the Late Cretaceous of Mongolia and its biological implications. Acta Palaeontol. Pol. 63, (2018).

59. Horner, J. R. & Padian, K. Age and growth dynamics of Tyrannosaurus rex. Proc. R. Soc. Lond. Ser. B: Biol. Sci. 271, 1875–1880 (2004).

60. Woodward, H. N. et al. Growing up Tyrannosaurus rex: Osteohistology refutes the pygmy “Nanotyrannus” and supports ontogenetic niche partitioning in juvenile Tyrannosaurus. Sci Adv 6, eaax6250 (2020).

61. Mallon, J. C., Bura, J. R., Schumann, D. & Currie, P. J. A Problematic Tyrannosaurid (Dinosauria: Theropoda) Skeleton and Its Implications for Tyrannosaurid Diversity in the Horseshoe Canyon Formation (Upper Cretaceous) of Alberta. Anat Rec 303, 673–690 (2020).

62. Stein, K. & Prondvai, E. Rethinking the nature of fibrolamellar bone: an integrative biological revision of sauropod plexiform bone formation. Biol. Rev. 89, 24–47 (2014).

63. Raguin, E., Rechav, K., Shahar, R. & Weiner, S. Focused ion beam-SEM 3D analysis of mineralized osteonal bone: lamellae and cement sheath structures. Acta Biomater 121, 497–513 (2021).

64. Currey, J. D. Bones: Structure and Mechanics. (Princeton University Press, Princeton, NJ, 2002).

65. Shapiro, F. & Wu, J. Woven bone overview: structural classification based on its integral role in developmental, repair and pathological bone formation throughout vertebrate groups. *Eur*. Cells Mater. 38, 137–167 (2019).

66. Ibrahim, J., Rechav, K., Boaretto, E. & Weiner, S. Three dimensional structures of the inner and outer pig petrous bone using FIB-SEM: Implications for development and ancient DNA preservation. J. Struct. Biol. 215, 107998 (2023).

67. Eltit-Guersetti, F., et al. Sclerotic prostate cancer bone metastasis: woven bone lesions with a twist. bioRxiv 2023.09.11.557266 (2023) doi:10.1101/2023.09.11.557266.

68. Weiner, S., Traub, W. & Wagner, H. D. Lamellar bone: Structure-function relations. J Struct Biol 126, 241–255 (1999).

69. Currey, J. D. Differences in the Blood-supply of Bone of Different Histological Types. J. Cell Sci. s3-101, 351–370 (1960).

70. Currie, P. J. & Koppelhus, E. B. Introduction to Albertosaurus Special Issue. Can. J. Earth Sci. 47, 1111–1114 (2010).

71. Coppock, C. C. & Currie, P. J. Additional Albertosaurus sarcophagus (Tyrannosauridae, Albertosaurinae) material from the Danek bonebed of Edmonton, Alberta, Canada with evidence of cannibalism. Can. J. Earth Sci. 61, 401–407 (2023).

72. Carr, T. D. A taxonomic assessment of the type series of Albertosaurus sarcophagus and the identity of Tyrannosauridae (Dinosauria, Coelurosauria) in the Albertosaurus bonebed from the Horseshoe Canyon Formation (CampanianMaastrichtian, Late Cretaceous). Can. J. Earth Sci. 47, 1213–1226 (2010).

73. Tanke, D. H. & Currie, P. J. A history of Albertosaurus discoveries in Alberta, Canada. Can. J. Earth Sci. 47, 1197–1211 (2010).

74. Osborn, H. F. Article XIV.-Tyrannosaurus and Other Cretaceous Carnivorous Dinosaurs. Proc. Acd. Nat. Sci. Phila 8, 72 (1905).

75. Eberth, D. A. & Kamo, S. L. High-precision U–Pb CA–ID–TIMS dating and chronostratigraphy of the dinosaur-rich Horseshoe Canyon Formation (Upper Cretaceous, Campanian–Maastrichtian), Red Deer River valley, Alberta, Canada. Can. J. Earth Sci. 57, 1220–1237 (2020).

76. Currie, P. J. & Eberth, D. A. On gregarious behavior in Albertosaurus. Can. J. Earth Sci. 47, 1277–1289 (2010).

77. Therrien, F., Zelenitsky, D. K., Voris, J. T. & Tanaka, K. Mandibular force profiles and tooth morphology in growth series of Albertosaurus sarcophagus and Gorgosaurus libratus (Tyrannosauridae: Albertosaurinae) provide evidence for an ontogenetic dietary shift in tyrannosaurids1. Can. J. Earth Sci. 58, 812–828 (2021).

78. Bell, P. R. Palaeopathological changes in a population of Albertosaurus sarcophagus from the Upper Cretaceous Horseshoe Canyon Formation of Alberta, CanadaThis article is one of a series of papers published in this Special Issue on the theme Albertosaurus. Can. J. Earth Sci. 47, 1263–1268 (2010).

79. Currie, P. J. Allometric growth in tyrannosaurids (Dinosauria: Theropoda) from the Upper Cretaceous of North America and Asia. Can. J. Earth Sci. 40, 651–665 (2003).

80. Erickson, G. M. et al. Gigantism and comparative life-history parameters of tyrannosaurid dinosaurs. Nature 430, 772–775 (2004).

81. Bodzioch, A. Idealized Model of Mineral Infillings in Bones of Fossil Freshwater Animals, on the Example of Late Triassic Metoposaurs from Krasiejów (Poland). Austin J Earth Sci. 2, (2015).

82. Frost, H. M. Micropetrosis. J. bone Jt. Surg. Am. Vol. 42-A, 144–50 (1960).

83. KINGSMILL, V. J. & BOYDE, A. Mineralisation density of human mandibular bone: quantitative backscattered electron image analysis. J. Anat. 192, 245–256 (1998).

84. Milovanovic, P. et al. Bone tissue aging affects mineralization of cement lines. Bone 110, 187–193 (2018).

85. Busse, B. et al. Decrease in the osteocyte lacunar density accompanied by hypermineralized lacunar occlusion reveals failure and delay of remodeling in aged human bone. Aging Cell 9, 1065–1075 (2010).

86. Shah, F. A. et al. Micrometer-Sized Magnesium Whitlockite Crystals in Micropetrosis of Bisphosphonate-Exposed Human Alveolar Bone. Nano Lett 17, 6210–6216 (2017).

87. Milovanovic, P. & Busse, B. Phenomenon of osteocyte lacunar mineralization: indicator of former osteocyte death and a novel marker of impaired bone quality? Endocr Connect 1, R70– R80 (2020).

88. Milovanovic, P. et al. The Formation of Calcified Nanospherites during Micropetrosis Represents a Unique Mineralization Mechanism in Aged Human Bone. Small 13, (2017).

89. Shah, F. A. The many facets of micropetrosis – Magnesium whitlockite deposition in bisphosphonate-exposed human alveolar bone with osteolytic metastasis. Micron 168, 103441 (2023).

90. Bell, L. S., Kayser, M. & Jones, C. The mineralized osteocyte: A living fossil. Am. J. Phys. Anthr. 137, 449–456 (2008).

91. Plotkin, L. I. & Bellido, T. Osteocytic signalling pathways as therapeutic targets for bone fragility. Nat. Rev. Endocrinol. 12, 593–605 (2016).

92. Marotti, G., Ferretti, M. & Palumbo, C. The problem of bone lamellation: An attempt to explain different proposed models. J. Morphol. 274, 543–550 (2013).

93. Rubanov, S. & Munroe, P. R. FIB-induced damage in silicon. J. Microsc. 214, 213–221 (2004).

94. Repp, F. et al. Coalignment of osteocyte canaliculi and collagen fibers in human osteonal bone. J Struct Biol 199, 177--186 (2017).

95. Loreille, O. et al. Ancient DNA analysis reveals divergence of the cave bear, Ursus spelaeus, and brown bear, Ursus arctos, lineages. Curr. Biol. 11, 200–203 (2001).

96. Bada, J. L., Wang, X. S. & Hamilton, H. Preservation of key biomolecules in the fossil record: current knowledge and future challenges. Philos. Trans. R. Soc. Lond. Ser. B: Biol. Sci. 354, 77–87 (1999).

97. Wyckoff, R. W. G., Wagner, E., Matter, P. & Doberenz, A. R. COLLAGEN IN FOSSIL BONE*. Proc. Natl. Acad. Sci. 50, 215–218 (1963).

98. Pawlicki, R. Studies of the fossil dinosaur bone in the scanning electron microscope. Z. fur Mikrosk.-Anat. Forsch. 89, 393–8 (1975).

99. Eberth, D. A. & Braman, D. R. A revised stratigraphy and depositional history for the Horseshoe Canyon Formation (Upper Cretaceous), southern Alberta plains. Can. J. Earth Sci. 49, 1053–1086 (2012).

100. Salamon, M., Tuross, N., Arensburg, B. & Weiner, S. Relatively well preserved DNA is present in the crystal aggregates of fossil bones. Proc. Natl. Acad. Sci. 102, 13783–13788 (2005).

101. Schmidt-Schultz, T. H. & Schultz, M. Bone protects proteins over thousands of years: Extraction, analysis, and interpretation of extracellular matrix proteins in archeological skeletal remains. Am. J. Phys. Anthr. 123, 30–39 (2004).

102. Lee, Y.-C. et al. Evidence of preserved collagen in an Early Jurassic sauropodomorph dinosaur revealed by synchrotron FTIR microspectroscopy. Nat. Commun. 8, 14220 (2017).

103. Orgel, J. P. R. O. et al. The in situ supermolecular structure of type I collagen. Structure 9, 1061–1069 (2001).

104. Meyers, P.-Y., C., Lin & Y., S. Biological materials: Structure and mechanical properties. Progress in materials science 53, 1–206 (2008).

105. Ibrahim, J. et al. FIB-SEM Study of Archaeological Human Petrous Bones: 3D Structures and Diagenesis. Minerals 14, 729 (2024).

106. Tomassini, R. L. et al. First osteohistological and histotaphonomic approach of Equus occidentalis Leidy, 1865 (Mammalia, Equidae) from the late Pleistocene of Rancho La Brea (California, USA). PLoS ONE 16, e0261915 (2021).

107. Huttenlocker, A. K. & Botha-Brink, J. Bone microstructure and the evolution of growth patterns in Permo-Triassic therocephalians (Amniota, Therapsida) of South Africa. PeerJ 2, e325 (2014).

108. Locke, M. Structure of long bones in mammals. J. Morphol. 262, 546–565 (2004).

109. Reznikov, N., Chase, H., Brumfeld, V., Shahar, R. & Weiner, S. The 3D structure of the collagen fibril network in human trabecular bone: Relation to trabecular organization. Bone 71, 189–195 (2015).

110. Zou, Z. et al. Three-dimensional structural interrelations between cells, extracellular matrix, and mineral in normally mineralizing avian leg tendon. Proc Natl Acad Sci USA 117, 14102– 14109 (2020).

111. Reznikov, N., Almany-Magal, R., Shahar, R. & Weiner, S. Three-dimensional imaging of collagen fibril organization in rat circumferential lamellar bone using a dual beam electron microscope reveals ordered and disordered sub-lamellar structures. Bone 52, 676–683 (2013).

112. Micheletti, C. et al. Encompassing the mesoscale in the multiscale characterization of osseointegration: A study of bone response to an additively manufactured implant for local genistein delivery. Sci. Rep. **Under review**, (2023).

113. Ayoubi, M. et al. 3D Interrelationship between Osteocyte Network and Forming Mineral during Human Bone Remodeling. Adv Healthc Mater 10, 2100113 (2021).

114. Morse, J. W. & Casey, W. H. Ostwald processes and mineral paragenesis in sediments. Am. J. Sci. 288, 537–560 (1988).

115. Monasterio-Guillot, L., Crespo-López, L., Navarro, A. B. R. & Álvarez-Lloret, P. Comparative Study of the Mineralogy and Chemistry Properties of Elephant Bones: Implications during Diagenesis Processes. Minerals 12, 1384 (2022).

116. Schindelin, J., et al. Fiji: An open-source platform for biological-image analysis. Nat Methods 9, 676–682 (2012).

117. Püspöki, Z., Storath, M., Sage, D. & Unser, M. Focus on Bio-Image Informatics. *Adv. Anat.*, Embryol. Cell Biol. 219, 69–93 (2016).

118. Fonck, E. et al. Effect of Aging on Elastin Functionality in Human Cerebral Arteries. Stroke 40, 2552–2556 (2009).

119. Rezakhaniha, R. et al. Experimental investigation of collagen waviness and orientation in the arterial adventitia using confocal laser scanning microscopy. Biomech. Model. Mechanobiol. 11, 461–473 (2012).

