## Supplemental Figure for "3D Focused Ion Beam Microscopy of Fossilized *Albertosaurus sarcophagus* reveals Nano to Microscale Structures"

[1.] School of Biomedical Engineering, McMaster University, Hamilton, Ontario, Canada
[2.] Fibics Incorporated, Ottawa, Ontario, Canada
[3.] Beaty Centre for Species Discovery and Palaeobiology Section, Canadian Museum of Nature, Ottawa, Ontario, Canada
[4.]Department of Earth Sciences, Carleton University, Ottawa, Ontario, Canada
[5.]Department of Materials Science and Engineering, McMaster University, Hamilton, Ontario, Canada
[6.] Canadian Centre for Electron Microscopy, McMaster University, Hamilton, ON, Canada

* Corresponding author:
Prof. Kathryn Grandfield
McMaster University
1280 Main Street West
Hamilton, L8S 4L7 Ontario, Canada


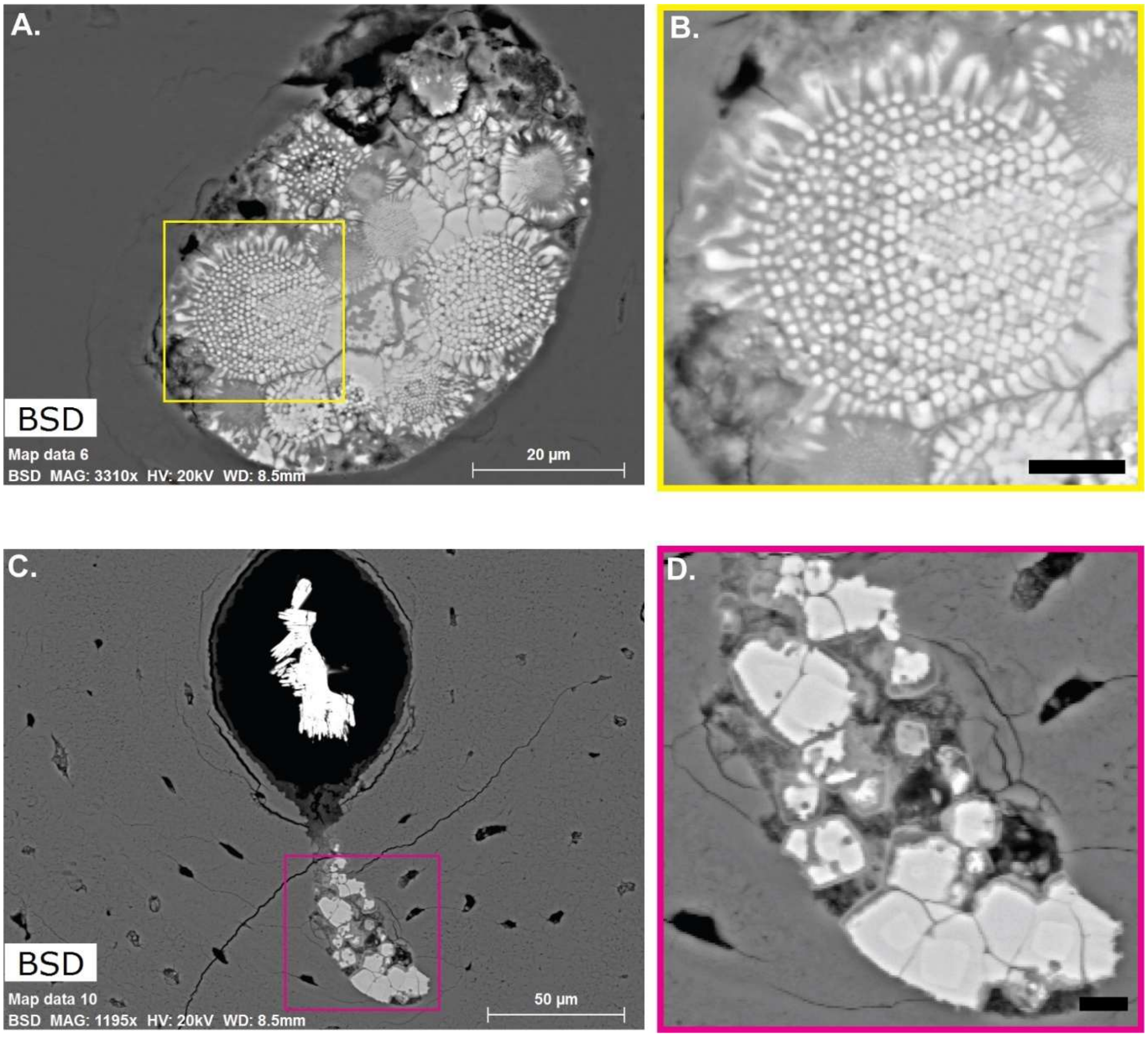


Figure S1. Comparison of pyrite structures. A) Backscatter image of smectite and framboidal pyrite infilled canal with the pyrite structures displaying the "sunflower" morphology. B) Inset from panel A) displaying a "sunflower" framboidal pyrite structure. C) Backscatter image of canal infilled with baryte crystals, smectite and recrystallized framboidal pyrite structures. D) Inset of panel C) displaying the recrystallized framboidal pyrite with areas of dissolution around the periphery and a layer of smectite coating the framboidal pyrite structure.

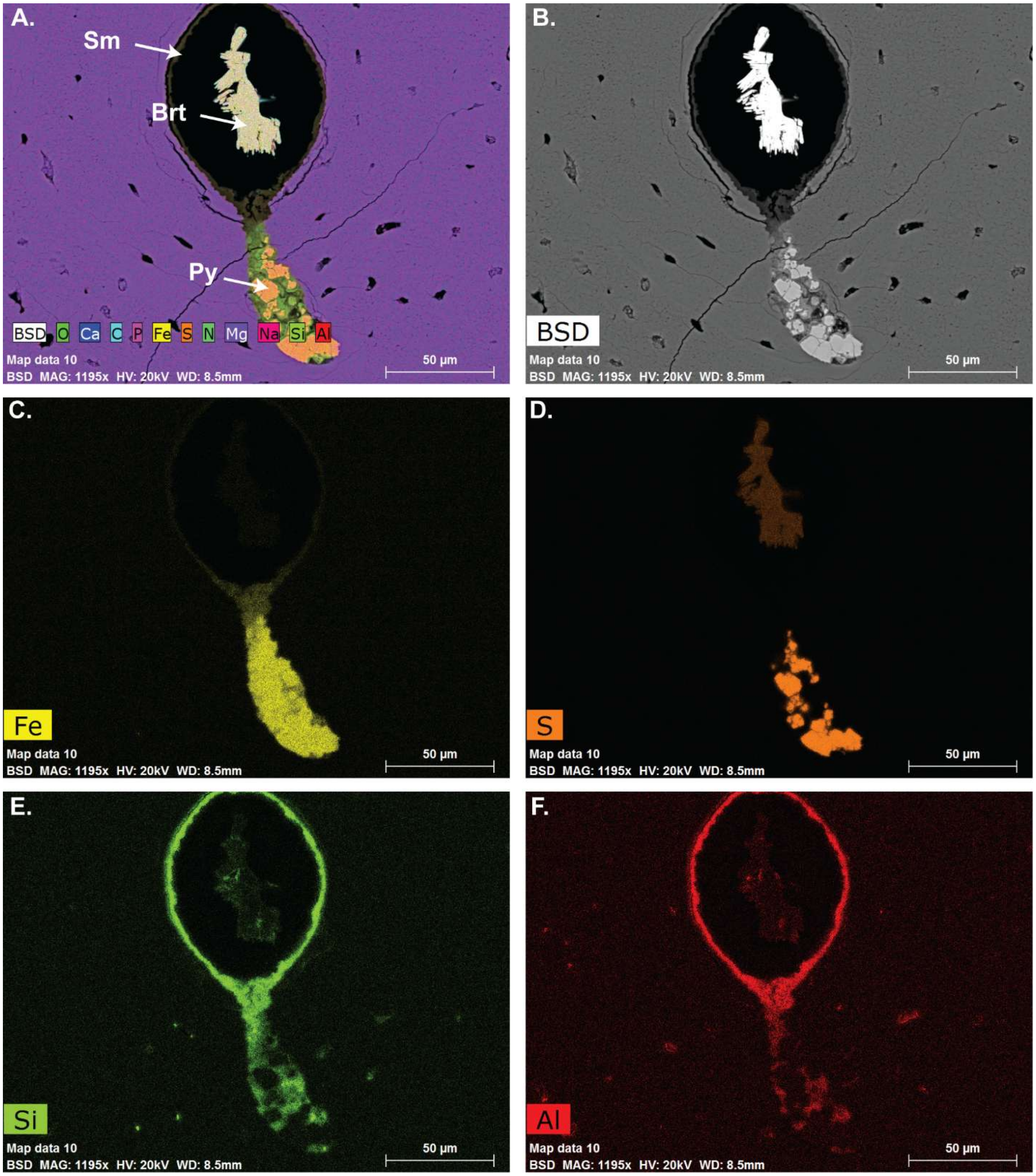


Figure S2. Element distribution maps showing the pyritization and silification in a Haversian canal and LCN mineral infiltration in the lateral region. A) EDS map overlay of all detected elements showing a cavity that is branching from the canal which was filled with secondary minerals, including pyrite (Py) and smectite-group minerals (Sm), due to diagenetic processes. The center of the Haversian canal is occupied by baryte crystals (Brt). B) Backscattered electron image of the haversian canal shown in A). Map of detected C) iron, D) sulphur, E) silicon, and F) aluminum. The cavity was filled with pyrite crystals (Py) and smectite-group (Sm) minerals. Sm minerals also line the Haversian canal and are also present in nearby LCN spaces based on the iron, silicon and aluminum element distribution maps.

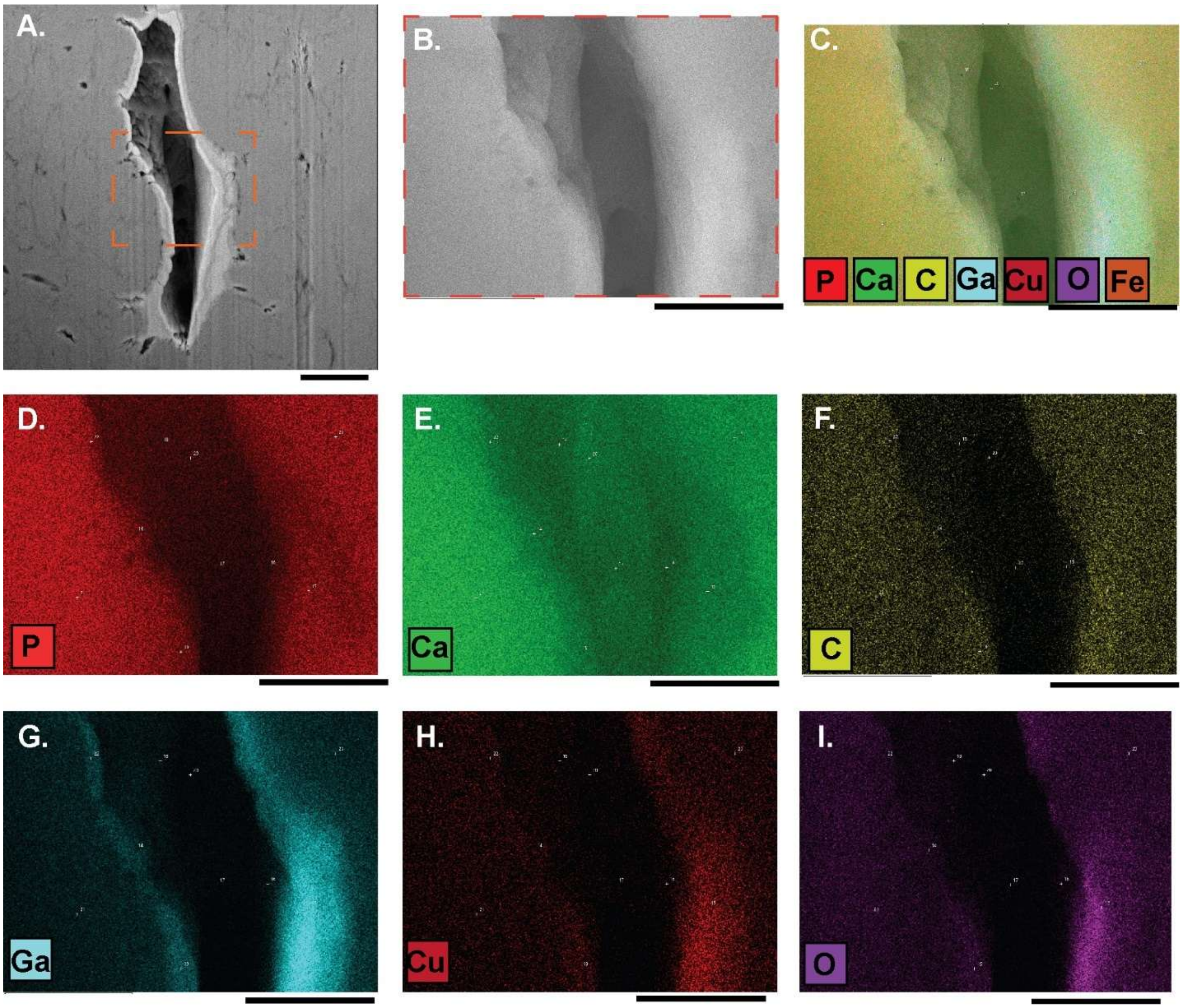


Fig. S3 Elemental maps of the cross-section of a lacuna in the lateral region (Fig 1D-Box 4). A) SEM image of the lacunae embedded in the bone matrix. B) Magnified SE2 image of the inside of the lacunae space. C) EDS overlay map of detected elements from the region in B. EDS map of detected D) phosphorous, E) calcium, F) carbon, G) gallium, H) oxygen and I) copper. Scale bars for A-I): 2.5 μm

Table S1. Imaging and Resolution details of Focused Ion Beam Scanning Electron Microscopy Acquisition Sites.

| | ***Run 1*** | ***Run 2*** | ***Run 3*** | ***Run 4*** |
|---|---|---|---|---|
| **Voxel size** | 20 nm | 10 nm | 10 nm | 10 nm |
| **SEM Imaging probe** | 2.0 kV; 60 µm | 2.0 kV; 30 µm | 2.0 kV; 30 µm | 2.0 kV; 30 µm |
| **Dwell x line average** | 8.0 µs x 5 | 3.7 µs x 4 | 6.0 µs x 6 | 3.6 µs x 4 |
| **EsB Grid Voltage** | 750 V | 750 V | 750 V | 750 V |
| **FIB Milling Probe** | 30 kV; 3 nA | 30 kV; 700 pA | 30 kV; 700 pA | 30 kV; 700 pA |
| **FIB Extractor Voltage** | -6.719 kV | -6.719 kV | -6.718 kV | -6.718 kV |

Table S2. Image Processing Steps for each FIB-SEM dataset performed within ORS Dragonfly after undergoing CLAHE contrast enhancement using FIJI.

| | | ***Run 1*** | ***Run 2*** | ***Run 3*** | ***Run 4*** |
|---|---|---|---|---|---|
| **FIB-SEM data signal** | | SE2 | SE2 | EsB | SE2 |
| **Shading Compensation (Polynomial)** | Degree | 5 | N/A | N/A | N/A |
| | Compensation | Subtract | N/A | N/A | N/A |
| **Slope Map** | Scale | N/A | 7 | N/A | N/A |
| **Vertical Destriping** | Sigma foreground | 128 | 128 | N/A | N/A |
| | Sigma background | 256 | 256 | N/A | N/A |
| | Level | 5 | 5 | N/A | N/A |
| | Wavelet | dbl2 | dbl2 | N/A | N/A |
| **Gaussian smoothing** | Level | 3D | N/A | 3D | N/A |
| | Size | 3 | N/A | 5 | N/A |
| | Standard deviation | 1.2 | N/A | 1.8 | N/A |

**VIDEO CAPTIONS**

Video S1. 3D rendered volume i) slicing through the SE2 dataset displaying parallel fibered bone, ii) rendering of collagen fibrils (green) and canaliculi (pink) in low mineralized regions that were visible and segmented using a U-Net deep learning segmentation, iii) A colour lookup table was applied to visualize individual mineral clusters [3D volume is represented as an orthographic projection and a scale bar is displayed]

Video S2. 3D rendered volume i) slicing through the SE2 dataset displaying parallel fibered bone near Volkmann's canal ii) rendering ofcollagen fibrils (green) and canaliculi (pink) in low mineralized regions that were visible and segmented using U-Net deep learning segmentation. The yellow arrow highlights collagen fibril bundles. [3D volume is represented as an orthographic projection and a scale bar is displayed]

Video S3. 3D rendered volume i) slicing through the EsB dataset of lamellar fibered bone near a primary osteon ii) rendering of collagen fibrils (green) and canaliculi (pink) in low mineralized regions that were visible and segmented using U-Net deep learning segmentation. [3D volume is represented as an orthographic projection and a scale bar is displayed]

Video S4. 3D rendered volume i) slicing through the SE2 dataset displaying a lacunocanalicular network ii) rendering of collagen fibrils (green) and canaliculi (pink) in low mineralized regions were visible and segmented using U-Net deep learning segmentation. [3D volume is represented as an orthographic projection and a scale bar is displayed]
